# Extreme storms cause rapid, context-dependent shifts in nearshore subtropical bacterial communities

**DOI:** 10.1101/801886

**Authors:** Ángela Ares, Margaret Mars Brisbin, Kirk N. Sato, Juan P. Martín, Yoshiteru Iinuma, Satoshi Mitarai

## Abstract

Climate change scenarios predict tropical cyclones will increase in both frequency and intensity, which will escalate the amount of terrestrial run-off and mechanical disruption affecting coastal ecosystems. Bacteria are key contributors to ecosystem functioning, but relatively little is known about how they respond to extreme storm events, particularly in nearshore subtropical regions. In this study, we combine field observations and mesocosm experiments to assess bacterial community dynamics and changes in physicochemical properties during early- and late-season tropical cyclones affecting Okinawa, Japan. Storms caused large and fast influxes of freshwater and terrestrial sediment—locally known as red soil pollution—and caused moderate increases of macronutrients—especially SiO_2_ and PO_43_^-^. We detected shifts in relative abundances of marine bacteria and the introduction of terrestrially-derived bacteria, including putative coral and human pathogens, during storm events. Soil input alone did not substantially affect marine bacterial communities in mesocosms, indicating that other components of run-off or other storm effects likely exert a larger influence on bacterial communities. The storm effects were short-lived and bacterial communities quickly recovered following both storm events. The early- and late-season storms caused different physicochemical and bacterial community changes, demonstrating the context-dependency of extreme storm responses in a subtropical coastal ecosystem.

## Introduction

Extreme storm events, such as tropical cyclones (i.e. tropical storms, hurricanes, and typhoons), can have dramatic consequences on coastal ecosystems, due in part to the effects of terrestrially-derived pollution (Hennessy *et al*., 1997; De Jesus Crespo *et al*., 2019). In addition to influencing salinity and turbidity, flood plumes often include elevated concentrations of bacteria (Solo-Gabriele *et al*., 2000), nutrients (i.e. C, N, P) (Chen *et al*., 2012, 2018; Gao *et al*., 2014; Paerl *et al*., 2018) and other chemicals, such as herbicides or heavy metals (Lewis *et al*., 2012; Mistri *et al*., 2019), which can act synergistically to negatively affect coastal ecosystems (Wooldridge, 2009; Brodie *et al*., 2012; Lewis *et al*., 2012). Especially in tropical and subtropical coastal ecosystems, which experience severe seasonal storms, large volumes of terrestrial run-off can enter coastal waters and can degrade coral reefs directly, through sedimentation or disease (Riegl and Branch, 1995; Philipp and Fabricius, 2003; Voss and Richardson, 2006; Haapkylä *et al*., 2011; Wilson *et al*., 2012). Such run-off events can also harm reefs indirectly, through eutrophication, hypoxia (Fabricius, 2005; Altieri *et al*., 2017) and decreased water quality. As global climate change is expected to enhance the frequency and intensity of extreme storm events (Groisman *et al*., 2005), it is increasingly important to better understand how such storms impact coastal ecosystem functioning.

Tropical cyclones are most active in the western North Pacific, where there is an average of 27 named storms per year (Wang *et al*., 2010; Herbeck *et al*., 2011) and where landfalling typhoons are specifically expected to become more common and more destructive (Mei and Xie, 2016). Okinawa Island, the largest island of the Ryukyu archipelago at the edge of the western North Pacific, is an ideal natural laboratory for studying storm effects on coastal ecosystems. Okinawa’s coral reefs have experienced significant declines in recent decades, due in part to increased storm induced run-off and sedimentation (Omori, 2011; Hongo and Yamano, 2013; Harii *et al*., 2014), which is exacerbated by agricultural practices and large coastal development projects (Omija, 2004; Masucci and Reimer, 2019). The fine-particle, laterite soils with high iron concentrations found in Okinawa and typical to the region are easily suspended and turn coastal waters a deep, cloudy red color during the frequent tropical cyclones (Supplemental Figure 1) (Omija, 2004). These events are locally referred to as Red Soil Pollution (Omori, 2011).

While the biological consequences of storm-induced run-off have been investigated for corals and fish species in Okinawa (Hongo and Yamano, 2013; Inoue *et al*., 2014; Yamazaki *et al*., 2015; O’Connor *et al*., 2016; Yamano and Watanabe, 2016), less is known about how tropical cyclones and the associated run-off affects coastal microbial communities and especially bacteria (Blanco *et al*., 2008). Microbial communities contribute to marine ecosystems through primary production and by recycling dissolved organic carbon and nutrients through the microbial loop (Azam *et al*., 1983), but can also draw down dissolved oxygen (Anderson and Taylor, 2001) and can cause opportunistic infections in marine organisms (Shinn *et al*., 2000; Sutherland *et al*., 2011; Sheridan *et al*., 2014; Peters, 2015). Therefore, changes in microbial community compositions in response to storms could precipitate large-scale ecosystem effects. Microbial responses can occur extremely quickly; Gammaproteobacteria, Flavobacteria and many Alphaproteopbacteria can increase in abundance within hours when exposed to high nutrient concentrations, whereas the entire microbial community—including archaea, protists, and viruses—can turn over on the scale of less than one day to about a week (Fuhrman *et al*., 2015). Rapid microbial response times to changing environmental conditions make microbes valuable early-warning bioindicators (Glasl *et al*., 2017; Pearman *et al*., 2018), but also hinders their study. Sampling at the scale of microbial response times during tropical cyclones is often dangerous and is further complicated by the poor predictability of storm tracks and event intensities (Zhou *et al*., 2012).

In this study, we characterize nearshore bacterial community dynamics in response to tropical cyclones affecting Okinawa Island and isolate the effects of sediment input through controlled mesocosm experiments. The study included tropical storm Gaemi at the start of the 2018 Okinawa typhoon season (June 16) and successive category 5 super typhoons, Trami and Kong-Rey, on September 30 and October 5, towards the end of the 2018 season. We evaluated physicochemical properties and bacterial community compositions in seawater samples collected before, during/between, and after storms in June and October and in samples taken from mesocosms with and without red soil amendment. Mesocosm experiments were undertaken in order to isolate the effects of red soil input from other storm effects, such as wind, waves, and fresh-water influx. The specific aims for this study were to: i) assess how bacterial community composition and physicochemical parameters respond in time to tropical cyclones and sediment input, ii) evaluate the speed of the responses and recovery, and iii) identify potential ecosystem consequences due to extreme storms and sediment input.

## Results

### Physicochemical responses to extreme storm events and red soil input

Two major storm events affecting Okinawa Island were monitored for this study. The first event included tropical storm Gaemi, which made landfall on June 16 and was the first tropical cyclone of the 2018 typhoon season. Gaemi brought 133.72 mm day^-1^ of precipitation and maximum wind speeds of 12.1 m s^-1^ to Okinawa (Figure 1). Substantial precipitation and wind were also recorded during the two days leading up to Gaemi’s landfall (69.77 and 83.99 mm day^-1^ precipitation and 10.2 and 12.1 m s^-1^ wind intensity on June 14 and 15, respectively), but no additional rain was recorded until July (Figure 1). The second event included two category-5 super-typhoons, Trami and Kong-Rey, which impacted Okinawa in rapid succession towards the end of the 2018 season (Sept. 29 and Oct.5). Trami and Kong-Rey caused accumulated rainfall of 239.24 and 87.89 mm day^-1^and maximum wind speeds of 26.9 and 12.7 m s^-1^, respectively. The June and October events represented the two largest rainfalls during the 2018 typhoon season, and typhoon Trami recorded the most extreme sustained and gusting wind speeds in 2018 (Figure 1). The extreme winds accompanying Trami and Kong-Rey also caused very large waves; Trami and Kong-Rey both brought waves with heights greater than nine meters (Japan Meteorological Agency, https://www.data.jma.go.jp/gmd/kaiyou/shindan/index_wave.html).

**Figure 1.**
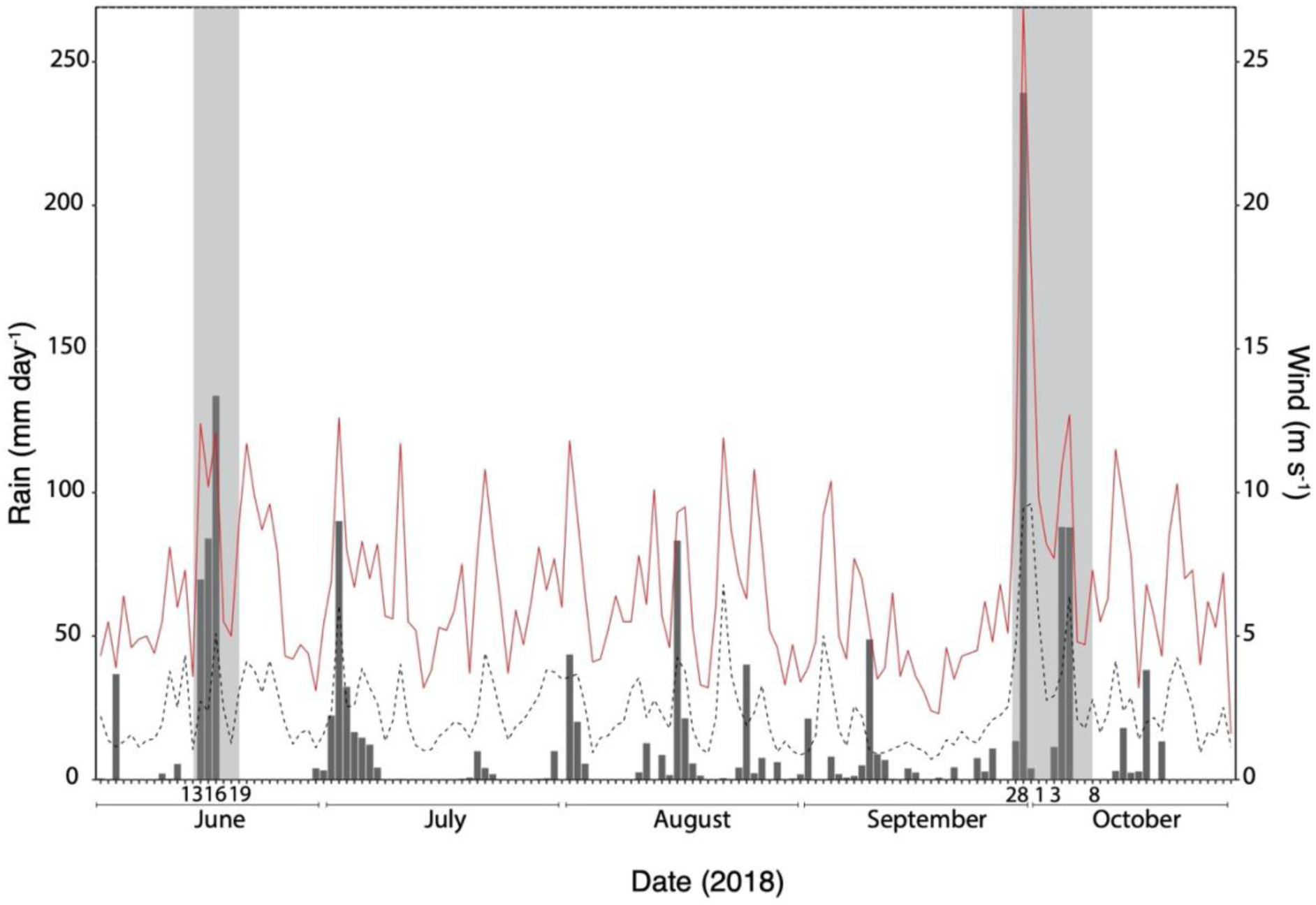
Precipitation (in mm day^-1^) and wind speed (m s^-1^) during the 2018 typhoon season in Okinawa, Japan. Data were collected with the meteorological station located at OIST Marine Science Station (26.510046 °N, 127.871721 °E) from June through October, corresponding to the duration of the typhoon season in Okinawa. Bars represent the daily amount of rain (in mm); dashed black and red lines indicate daily mean and maximum wind speeds (in m s-1); shaded areas represent the two red soil pollution events monitored in this study: Tropical Storm Gaemi on June 16 and the super-typhoons Trami and Kong-Rey on September 29 and October 5. The dates when water samples were collected for chemical and DNA analyses are noted on the x-axis.

Temperature (°C), DO (mg L^-1^), salinity (‰) and turbidity (NTU) were measured *in situ* at the four nearshore field sites at the same time that water samples were collected. There was a significant decrease in salinity (∼15‰) and concurrent increase of turbidity (∼10-fold) during the storm on June 16 compared to before the storm on June 13 and afterwards on June 19 (Figure 2). Up to a 5‰ decrease in salinity and 3 NTU increase in turbidity was measured between the two storms in October (Oct 1 and 3). These changes were smaller in magnitude than were recorded for the June storm and were not statistically significant, despite the storms in October delivering much more precipitation than the June storm (Figure 1). However, the wind accompanying the October storms was also much more intense (Figure 1), which presumably mixed the water column more thoroughly and prevented freshwater lenses from forming, thus contributing to diminished changes in salinity and turbidity being observed in October compared to June.

**Figure 2.**
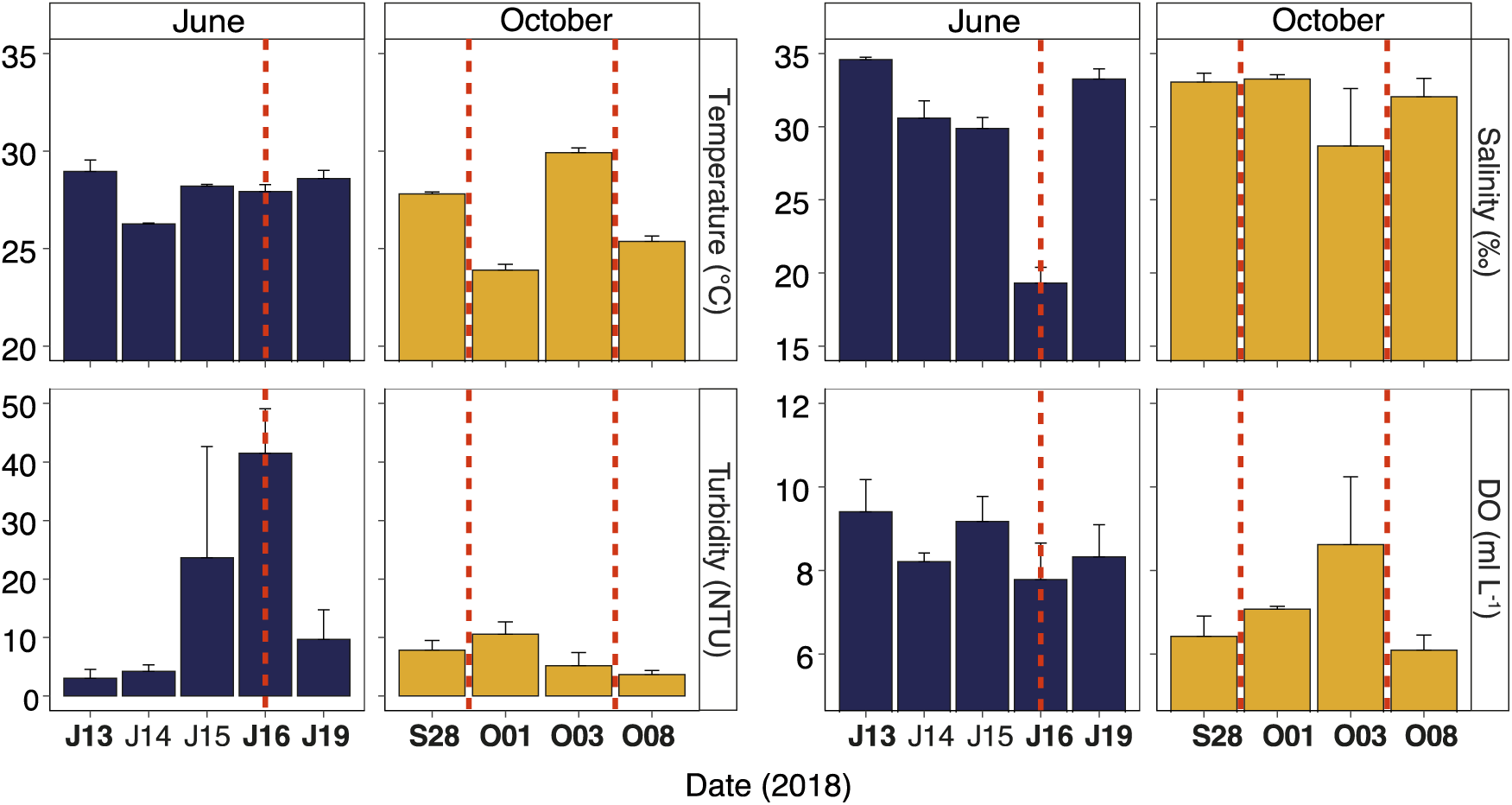
Temporal variation of temperature (°C), turbidity (NTU), salinity (‰) and dissolved oxygen (DO; mg L^-1^) before, during, and after storm events in June and October, 2018. Bars represent mean concentrations of dissolved temperature (°C), turbidity (NTU), salinity (‰) and DO (mg L^-1^) for four sampling sites along the central west coast of Okinawa, Japan. Sampling dates when samples were also processed for metabarcode analyses are indicated in bold on the x-axis. Red dashed vertical lines represent the timing of storms and corresponding red soil pollution events. All measurements were taken with a CTD profiler.

Concentrations of dissolved nutrients, including NO_2_^-^, NO_3_^-^, NH^4+^, PO_43_^-^, SiO_2_, and dFe, were measured in seawater samples collected in June and October (Figure 3) and throughout the second mesocosm experiment (Supplemental Figure 7). Overall, dissolved nutrient concentrations were higher in June than October, with the exception of dFe and SiO_2_, which both had similar values in the two sampling months (Figure 3). A nonparametric Kruskall-Wallis test was applied to detect significant storm-induced differences in field nutrient concentrations (Supplemental Table 3). Results showed significant increases (*p* < 0.05) in NO_2_^-^ concentrations during and following storm events in June and October, whereas SiO_2_ and PO_43_^-^ were only significantly elevated during and after the storm in June (Figure 3).

**Figure 3.**
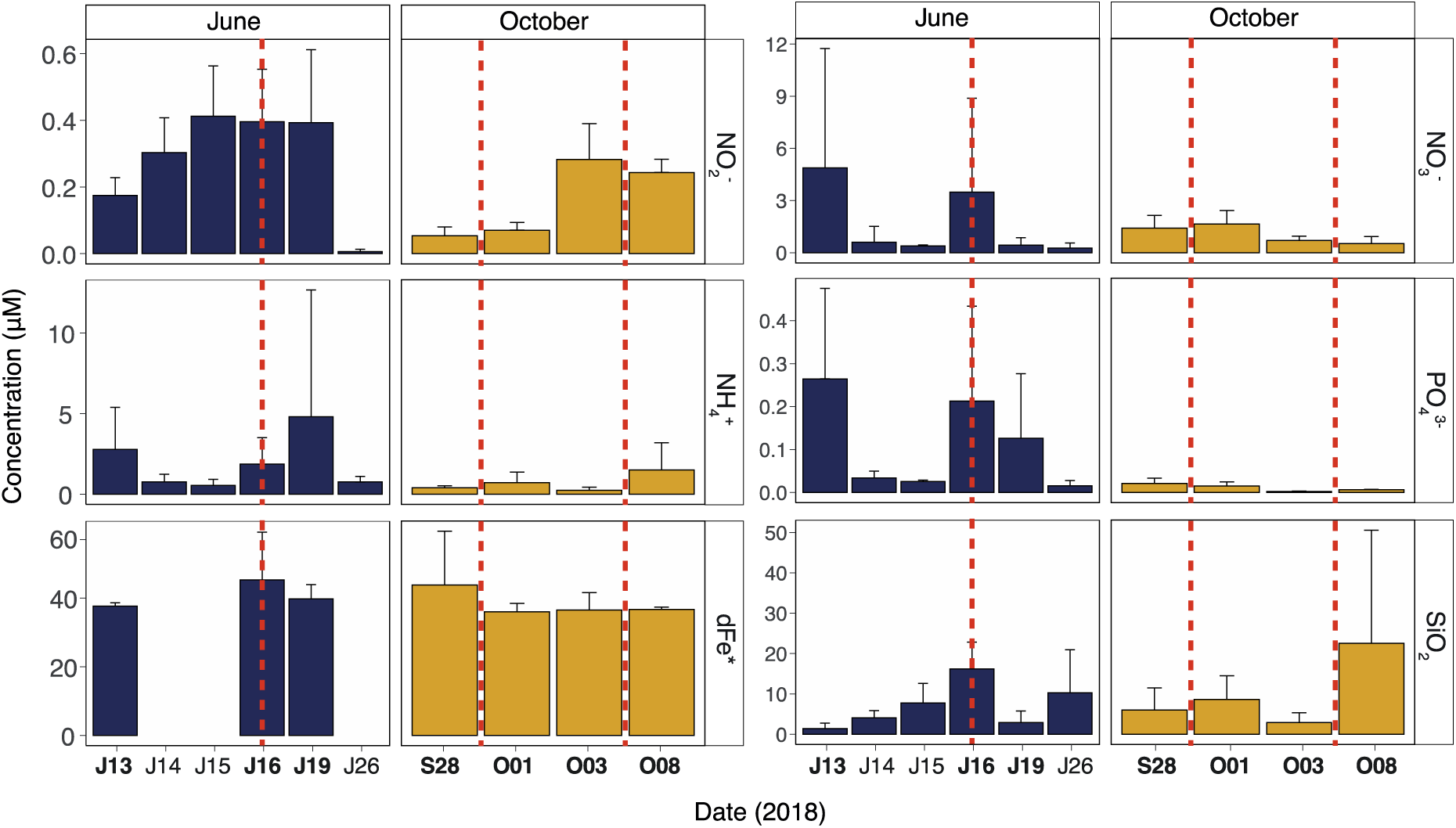
Temporal variation of micro- and macro-nutrient concentrations before, during, and after storm events in June and October, 2018. Bars represent mean concentrations of dissolved NO_2_^-^, NO_3_^-^, NH_4_^+^, PO_43_^-^, SiO_2_, and dFe (* dFe concentration is in nM) for four sampling sites along the central west coast of Okinawa, Japan. Error bars represent one standard deviation of the mean from four replicates. Red dashed vertical lines represent the timing of major storms and associated red soil pollution events. Sampling dates when samples were also processed for metabarcode analyses are indicated in bold on the x-axis. Nutrient concentrations were determined on a QuAAtro39 Continuous Segmented Flow Analyzer and dFe concentration was determined by ICP-MS after Mg(OH)^2^ co-precipitation using the isotope dilution method (Wu and Boyle, 1998).

Red soil addition in the October mesocosm experiment caused a significant increase in SiO_2_ concentration (Supplemental Figure 7, Supplemental Table 4). Additionally, red soil addition caused PO_43_^-^ concentrations to increase above the detection limit 4 hours following soil addition, whereas PO_43_^-^ stayed below the detection limit in control mesocosms after the initial measurement (Supplemental Figure 7). Two-way ANOVA results (Supplemental Table 4) indicate that time had a greater effect on nutrient concentration (*p* < 0.05 for NO_2_^-^, SiO_2_ and dFe) than red soil treatment (*p* < 0.05 for SiO2 and dFe). The treatment by time interaction was only significant in the case of SiO_2_ (*p* < 0.05), for which higher concentrations were found in soil-treated mesocosms.

### Bacterial community responses to storm events and red soil input

Metabarcoding with the bacterial 16S ribosomal RNA gene was performed to evaluate shifts in bacterial community composition associated with extreme storms in the field and with sediment input in mesocosms. There was a clear shift in the relative abundances of bacterial phyla in field samples collected during the June storm compared to before or afterwards. Several phyla that were also present in soil samples became more abundant in the water samples collected during the storm—including Acidobacteria, Actinobacteria, Chloroflexi, Firmicutes, Rokubacteria and Verrumicrobia—but phyla absent in soil samples also became more abundant during the storm, most notably Epsilonbacteraeota (Figure 4). Principal coordinate analysis (PCoA) of Aitchison distances between bacterial community compositions in water samples also illustrated a clear shift in community composition during the June storm event; samples collected during the storm clustered separately along the primary axis from the samples collected both before and after the event (Figure 5). The shift in community composition during the storm was accompanied by an increase in ASV richness (Figure 6); estimated ASV richness was significantly higher in samples collected during the June storm compared to samples collected before and afterwards.

**Figure 4.**
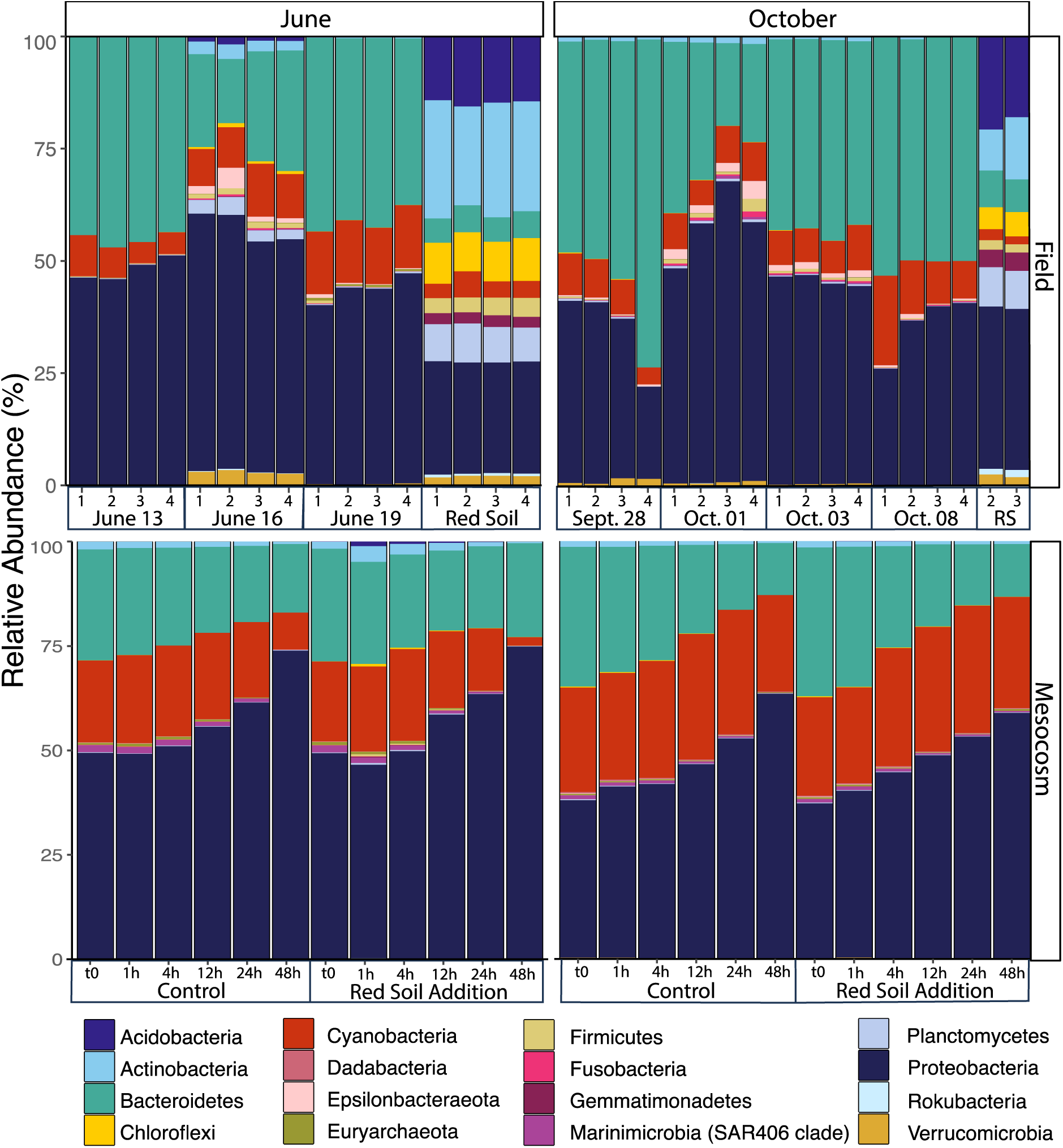
Relative abundance of prokaryotic phyla in field and mesocosm samples collected in June and October, 2018. Each stacked bar represents the relative contribution of major bacterial phyla to the total community at one sampling location, in one red soil sample, or at one time point in mesocosms (mesocosm bars represent aggregate data from 4–5 replicates depending on month and treatment). Sampling stations 1–4 were sampled before, during, and after red soil pollution events triggered by major storms in June and October 2018. In June, 6/13 was before the event, 6/16 was during the event, and the 6/19 was after the event. In October, 9/28 was before the event, 10/01 and 10/03 were between events, and 10/08 was after events. Red soil samples are labeled with replicate number and were collected close to the dates storms occured in June and October. Red soil (200 mg L^-1^) was added to treatment mesocosms immediately following the t0 sampling.

**Figure 5.**
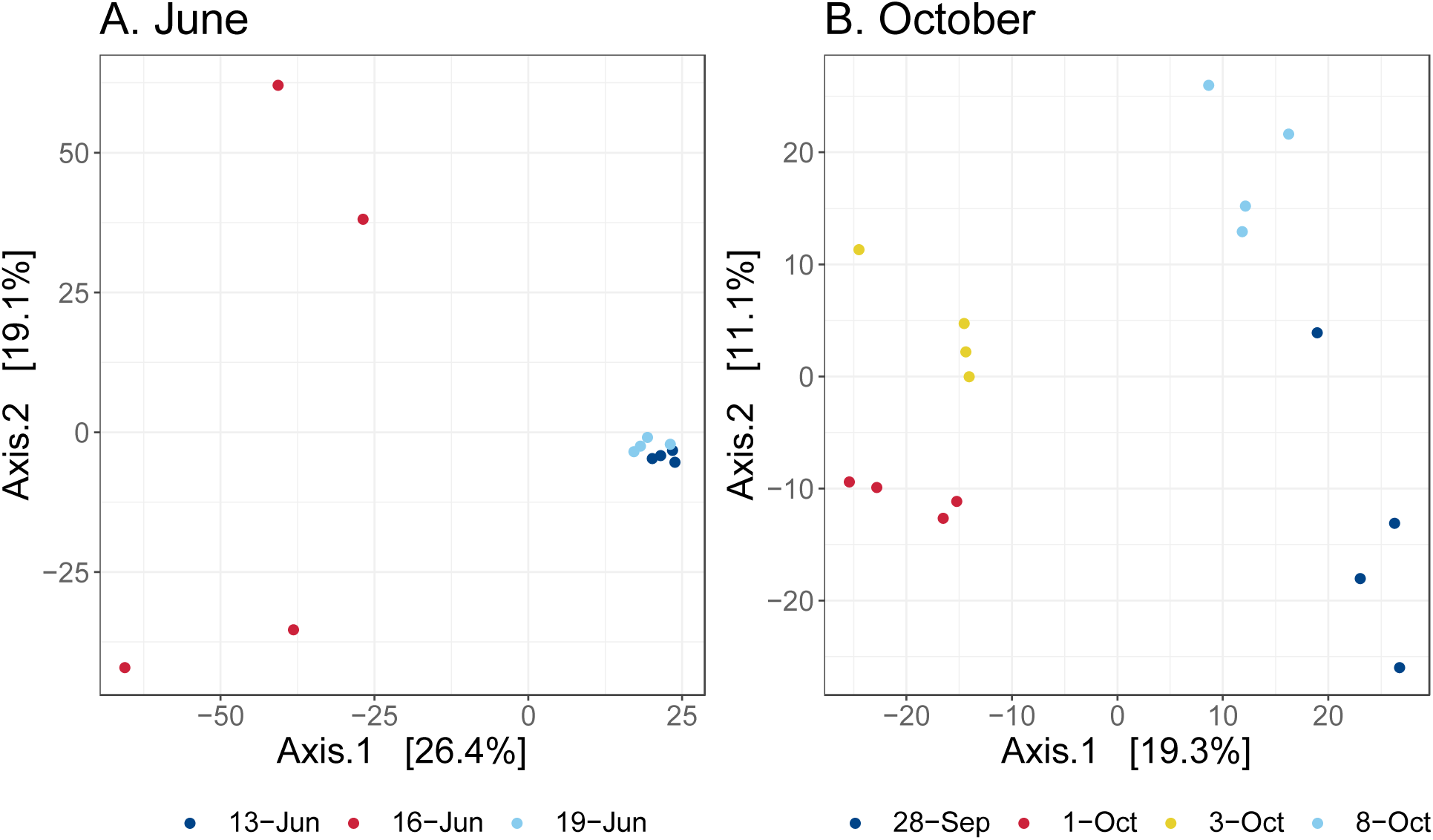
Principal coordinates analysis (PCoA) of Aitchison distances between prokaryotic community composition before, during, and after storm events in June (A) and October (B), 2018. (A) June 13 was before the June storm, 6/16 was during, and 6/19 was after. For the October event (B), 9/28 was before the storms, 10/01 and 10/03 were between, and 10/08 was after. Samples collected during/between storm events cluster separately from samples collected before and after for both the June and October events.

**Figure 6.**
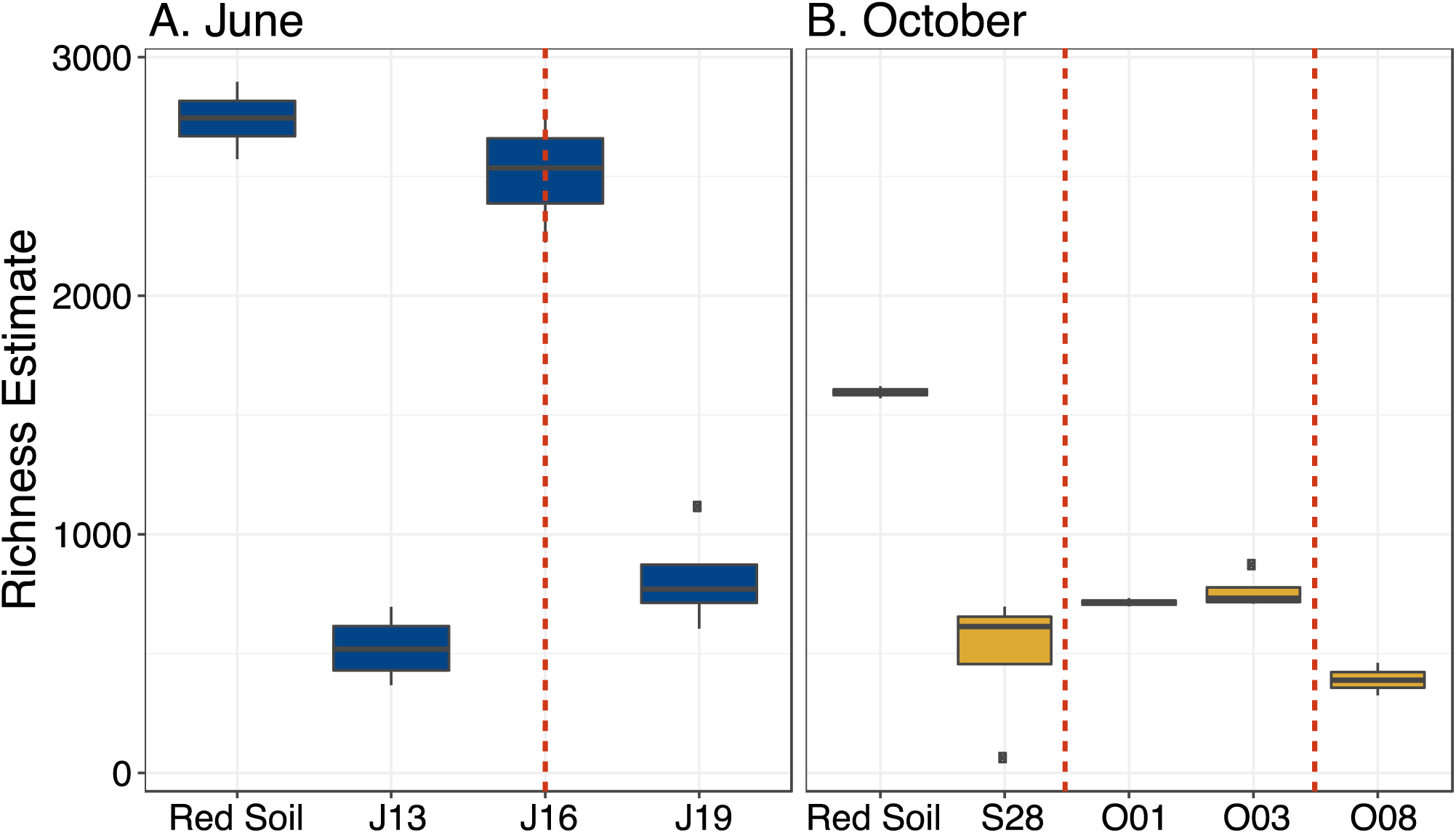
Richness estimates for bacterial communities in red soil samples and surface water samples collected before, during, and after storm events in June (A) and October (B), 2018. Red dashed vertical lines represent the timing of major storms and associated red soil pollution events. In June (A), the red soil samples had significantly higher richness than the water samples collected before and after the event, but not samples collected during the event. In October (B), red soil samples had significantly higher richness than all water samples and richness was not significantly different in water samples collected before, during, or after the storms. Differences in richness were considered significant when p < 0.05.

Soil addition to June mesocosms also influenced bacterial community composition and richness, although to a lesser extent than the storm-influenced nearshore communities. Bacterial phyla dominant in soil samples were detectable in mesocosm samples taken one hour after soil addition, but their abundance diminished in samples taken 24 hours later (Figure 5). Likewise, ASV richness increased in samples taken from mesocosms following soil addition and decreased over time (Supplemental Figure 9). Overall, mesocosm incubation time had a larger effect on bacterial community composition than soil addition (Figure 4, Supplemental Figure 10), despite mesocosm conditions (temperature, salinity, DO) remaining similar to ambient conditions throughout the experiment (Supplemental Figures 5–6).

In October, the community composition also shifted between the two storms relative to before and after the overall storm event. Additionally, some phyla present in soil samples, particularly Firmicutes, increased in relative abundance between the storms, but similar to in June, phyla absent from soil samples also increased in relative abundance (e.g. Epsilonbacteraeota and Fusobacteria). However, unlike in June, Actinobacteria, Planctomycetes, and Verrucomicrobia were already detectable in water samples collected before the storm event occurred (Figure 4). PCoA further demonstrated a shift in community composition in samples collected between the October storms compared to before and afterwards; samples collected between the two storms clustered separately on the primary axis from samples collected before and after the event (Figure 5). Interestingly, the estimated ASV richness was lower (by about half) for all samples collected in October compared to in June, including in soil samples, and estimated richness did not increase in the samples collected between storms (Figure 6). Community compositions in field samples from the different months segregated into two clusters along the primary axis in PCoA and the differences were statistically significant in PERMANOVA (*p* < 0.01, *F* = 5.37, *R*_2_ = 0.17), demonstrating that nearshore bacterial communities were distinct in June and October (Supplemental Figure 8). Furthermore, soil addition to October mesocosms did not cause observable increases in relative abundances of phyla found in soil samples as it had in June (Figure 4, Supplemental Figure 9).

In order to identify individual taxa that became more abundant during storms and, thus, contributed to the broader patterns observed in the data, we performed pairwise testing for differentially abundant ASVs between sampling dates in each month. The greatest number of significantly differentially abundant ASVs were found in the pairwise test between June 13 (before the storm) and June 16 (during the storm), with the vast majority being more abundant during the storm (Figure 7A). In contrast, very few ASVs were significantly differentially abundant in samples collected before and after the June storm (Figure 7A). While many of the ASVs that were significantly more abundant during the storm on June 16 were also detected in soil samples, the majority were not (Figure 7A, B). ASVs that were significantly more abundant on June 16 compared to June 13 belonged to a total of 67 orders—including Flavobacteriales, Campylobacterales, and Vibrionales, which can be pathogenic to humans and marine organisms (Pruzzo *et al*., 2005; Silva *et al*., 2011; Loch and Faisal, 2015; Canty *et al*., 2020), and Rhizobiales, Sphingomonadales, and Alteromonadales, which were also found in soil samples (Figure 7B). In pairwise tests for October samples, the largest number of differentially abundant ASVs were likewise found between samples collected before (Sept. 28) and during storms (Oct. 1 and 3), but none of the differentially abundant ASVs were also found in soil samples (Figure 7C, D). ASVs that were significantly differentially abundant between samples taken before and during the October storms belonged to 21 different orders, with most, but not all, also represented in the June results (Figure 7D).

**Figure 7.**
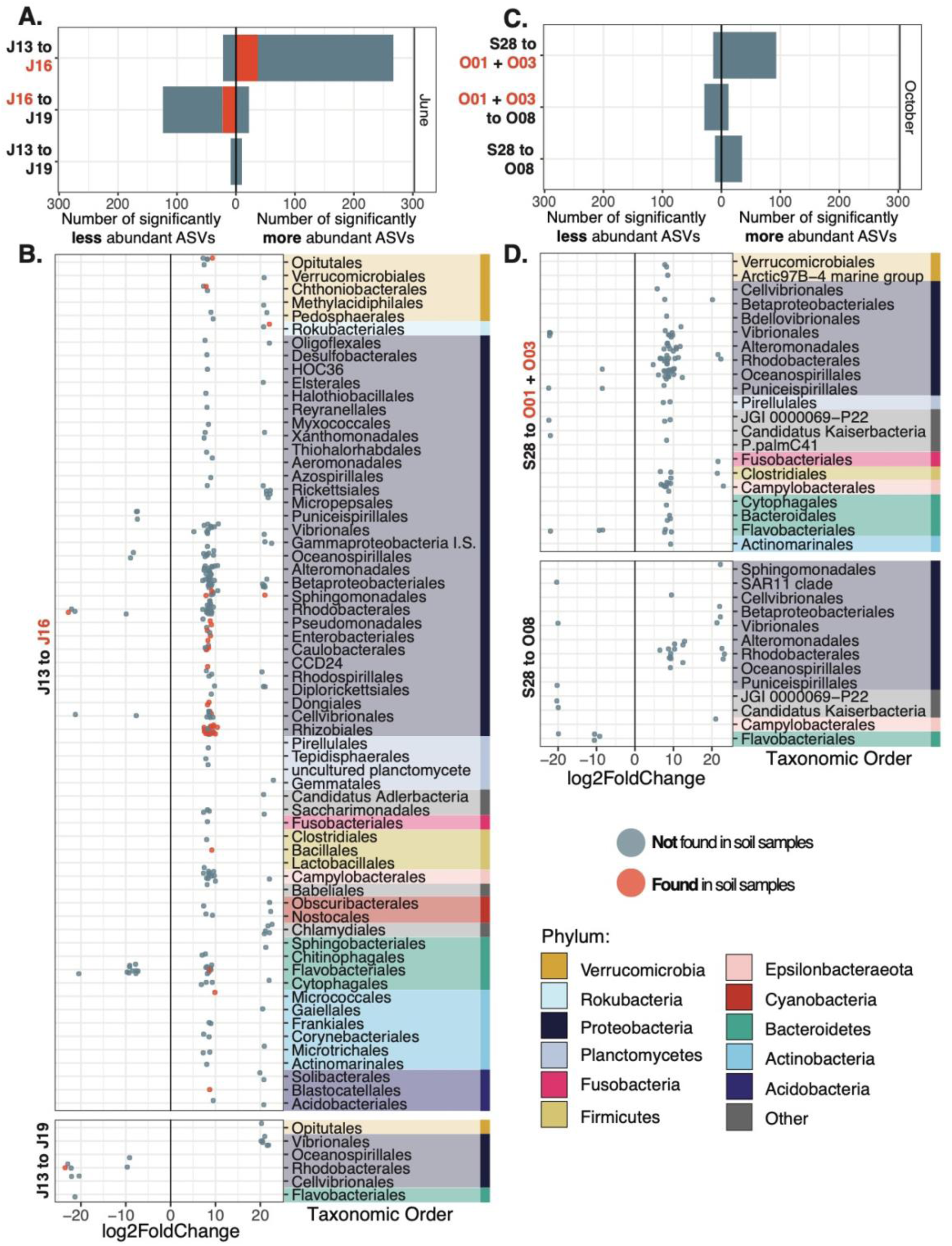
Number of Amplicon Sequence Variants (ASVs) significantly differentially abundant in pairwise comparisons between sampling dates in June (A) and October (C) and log2 Fold Change of individual ASVs in pairwise comparisons from before to during storms and from before to after storms in June (B) and October (D), 2018. ASVs were considered significantly differentially abundant when the FDR adjusted p-value was less than 0.05. ASVs that were also present in soil samples are colored red in panels A and B; no differentially abundant ASVs in pairwise tests for October samples were found to be present in October soil samples. Positive log2FoldChange values (x-axis, B, D) indicate higher abundance of ASVs during/between storms compared to before (J13 to J16, S28 to O01 + O03) or after storms compared to before (J13 to J19, S28 to O08). Differentially abundant ASVs are grouped by taxonomic order and orders are color coded by phylum so that colors correspond to Figure 4.

## Discussion

As the severity and frequency of extreme storm events increases with global climate change, it is increasingly important to understand how these events impact ecological functioning in marine ecosystems (Wetz and Paerl, 2008; Du *et al*., 2019). However, characterizing the effects of extreme storm events, such as typhoons, on coastal ecosystems is a complicated task, due both to forecast unpredictability, which makes sampling before and after events challenging, and the dangerous conditions that accompany storms (Chen *et al*., 2018). This study reports on the near-shore microbial community dynamics and relevant environmental parameters in two short time-series encompassing major storms during the 2018 typhoon season in Okinawa, Japan. In addition, concurrent, controlled mesocosm experiments were performed to supplement field observations and isolate impacts of terrestrial sediment input that regularly accompanies large storms in Okinawa. Predictably, storms caused influxes of both freshwater and sediment into the coastal marine environment (Figure 2, Supplemental Figure 1), which carried some soil-derived bacteria and some bacteria presumably derived from other terrestrial sources (Figure 4, Figure 7). Remarkably though, the effects of the storms were extremely short-lived, and bacterial community compositions quickly recovered. While field samples were collected three days following storm events, mesocosm experiments showed that the bacterial community began recovering only four hours after soil addition and was fully recovered after just 24 hours. Despite the rapidity of community shifts, terrestrially derived bacteria and marine bacteria that increased in abundance during and after storms can still be detrimental to coastal ecosystems and dangerous to human health. For instance, bacteria that became more abundant following storms can cause diseases in corals (e.g. *Vibrio spp*., *Pseudoalteromonas sp*., Rhodobacteraceae bacteria) and humans (e.g. Campylobacter, Fusobacteria, Enterobacter), thus necessitating a better understanding of the sources and sinks for these bacteria in the coastal environment (Pruzzo *et al*., 2005; Vizcaino *et al*., 2010; Silva *et al*., 2011; Zimmer *et al*., 2014; Figure 7).

### Extreme storms cause rapid changes in environmental conditions and bacterial community composition

During the storm event in June, we measured a drastic drop in salinity and increased turbidity in nearshore surface waters (Figure 2), which is similar to previous reports following major storm events (De Carlo *et al*., 2007; Zhou *et al*., 2012; Chen *et al*., 2018). The influx of freshwater and sediment was accompanied by moderate increases in NO_2_^-^, NO_3_^-^, NH^4+^, PO_43_^-^, SiO_2_ and dFe concentrations (Figure 3), which is consistent with terrestrial run-off from agricultural areas with high iron-content soil, as is found in Okinawa (Arakaki *et al*., 2005). However, increased nutrient loading dissipated before samples were collected again 3 days after the June storm (Figure 3). Likewise, there was an increase in bacterial diversity (Figure 6) representing the introduction of bacterial phyla that were found in our soil samples and are common components of soil microbiomes, including Acidobacteria, Actinobacteria, Chloroflexi, Firmicutes, Planctomycetes, Rokubacteria and Verrucomicrobia (Figure 4; Witt *et al*., 2012; Balmonte *et al*., 2016; Lin *et al*., 2019) and additional phyla presumably deriving from other marine or terrestrial sources, such as Epsilonbacteraeota. Three days following the storm, on June 19, the alpha diversity returned to pre-storm levels (Figure 6A) and the community composition was almost identical to before the storm (Figure 5A). Moreover, only 19 ASVs were significantly differentially abundant between June 13 and June 19 compared to 267 ASVs between June 13 and June 16 (Figure 7A), demonstrating the incredible speed at which the microbial community recovered. The ephemeral presence of bacteria also found in the soil samples was similarly apparent in the June mesocosm experiment. One hour after soil was added to treatment mesocosms, phyla dominant in soil samples (e.g. Acidobacteria, Chloroflexi, Firmicutes, and Planctomycetes) were detected, but their relative abundance decreased in samples taken four and twelve hours later, and they were absent in samples taken 24 hours later. The transience of storm-effects on nearshore bacterial communities was further demonstrated in results from the October storm event. The bacterial community recovered just three days after super-typhoon Kong-Rey passed, the second super-typhoon to affect Okinawa in less than a week (Figure 5B).

### Extreme storms cause context-dependent changes in environmental conditions and bacterial community composition

While storm effects were extremely transient in both cases (Figure 5), the June and October storm events affected coastal physicochemistry and bacterial community composition differently (Figure 2, 3 and 4). Events in both June and October increased coastal NO_2_^-^ loading, but only the June event was accompanied by increased PO_43_^-^ and SiO_2_ concentrations, and the June storm caused more pronounced shifts in bacterial community composition, highlighted by a larger increase in ASV richness and a higher number of differentially abundant ASVs. Although these differences may reflect disparate sampling schemes, it is likely that differences in wind and rain intensity and the contexts in which the two events occurred also had a strong effect. Key differences between the June and October events were that much more rain, wind and wave action accompanied the October storm event than the June storm and that the June storm made landfall at the beginning of the typhoon season, whereas the October storms affected Okinawa towards the end of the typhoon season.

Typhoon Trami, on Sept. 29, caused twice as much precipitation as tropical storm Gaemi, on June 16 (Figure 1), which could have diluted nutrient loading in storm run-off and caused the smaller changes in nutrient concentrations recorded in October compared to in June (Figure 3). When flushing rate is high, less PO_43_^-^ is desorbed from sediments and there is a dilution effect for both dissolved phosphorus and nitrogen in run-off (Blanco *et al*., 2010). Additionally, Gaemi was the first major storm of the 2018 typhoon season (June–October) following a relatively prolonged dry period, which could increase nutrient loading in storm run-off, especially since agricultural fertilizers are applied throughout the preceding spring and summer growing season. By October, several tropical storms and typhoons had already affected Okinawa (Figure 1); the 2018 Pacific typhoon season had higher than average storm frequency and included 29 tropical storms, 13 typhoons, and 7 super typhoons, although not all made direct landfall with Okinawa (Japan Meteorological Agency, https://www.jma.go.jp/jma/indexe.html). These intervening events could have stripped topsoil and depleted soil nutrients and microbiomes, so that October storm run-off carried less nutrients, organic material, and terrestrial bacteria into coastal water than storm run-off in June. Moreover, antecedent soil moisture affects dissolved nutrient loading in run-off, with less nutrients desorbing from clay-based soils, like Okinawa red soil, when wet (Perrone and Madramootoo, 1998).

The controlled mesocosm experiments offer additional insight for interpreting differences in field observations between June and October. Despite collecting soil from the same place in June and October, the soil microbiome was much less diverse in October (Figure 5) and soil addition to October mesocosms did not introduce soil bacteria or increase bacterial diversity as it had in June mesocosms (Figure 4, Supplemental Figure 9). These results suggest that soils in Okinawa could have lower bacterial loading towards the end of the typhoon season. Furthermore, soil addition in October mesocosms did not cause nitrogen (NO_2_^-^ or NO_3_^-^) or dFe to increase over baseline measurements, but did cause increases in SiO_2_ and PO_43_^-^ (Supplemental Figure 7). However, the increase in SiO_2_ and PO_43_^-^ were gradual, demonstrating that time is required to release these compounds from red soil (De Carlo *et al*., 2007; Blanco *et al*., 2010; Chen *et al*., 2018). Soils in Okinawa could, therefore, have lower nitrogen content at the end of the typhoon season and more intense rains may deliver less nutrients due to rapid flushing.

### Influence of extreme storms on bacterial taxa in nearshore waters

The influence of June and October storms on coastal microbial communities varied in magnitude, but in both instances the community shift was rapid and transient and included several common bacterial groups that became more abundant; 15 out of the 21 bacterial orders encompassing ASVs that were significantly more abundant during the October event were shared with the June event (Figure 7). These shared bacterial orders included primarily heterotrophic bacteria, several of which contain potentially pathogenic groups (e.g. Vibrionales, Fusobacteriales, and Campylobacterales) (Pruzzo et al. 2005; Silva et al. 2011; Canty et al. 2020). Originally, we expected cyanobacteria to respond to increased inorganic nutrients delivered to the coastal ecosystem with storm run-off. Specifically, we expected nitrogen fixers—such as *Trichodesmium*, which occasionally blooms near Okinawa (Grossmann *et al*., 2015)—to benefit from increased dFe and PO_43_^-^ in run-off (Sañudo-Wilhelmy *et al*., 2001; Wu *et al*., 2003). Instead, only a few cyanobacteria ASVs became more abundant during the June storm, but not the October storm (Figure 7). Namely, cyanobacteria belonging to the family Oscillatoriaceae, which form filamentous benthic mats (Siegesmund *et al*., 2008; Engene *et al*., 2018), and Obscuribacterales, which are uncultured and have unknown morphology, but may not be photosynthetic (Soo *et al*., 2014), became more abundant in June. There are several possible explanations for the minimal changes in cyanobacteria relative abundance: i) cyanobacteria were not limited by compounds present in run-off, ii) nutrients in run-off were not biologically available, or iii) terrestrial run-off was transported offshore or diluted too quickly to affect nearshore communities. Nutrient concentrations that were elevated during the June storm (NO_2_^-^, NO_3_^-^, PO_43_^-^ and dFe) decreased after the storm passed, but not immediately and not as quickly as the microbial communities recovered (Figure 3). Since nutrients were not immediately drawn down, the nearshore microbial community may not have been nutrient limited in our study area. The limited bacterial response to soil addition in mesocosms, despite increased PO43- and dFe in October, further suggests that nutrient availability was not driving community composition. The heterotrophic bacteria that became more abundant during and after storms could have derived from the soil or other components of the run-off, been resuspended from the benthos due to wind and waves, or increased in abundance in response to organic matter in the run-off.

### Potential ecosystem consequences due to terrestrial run-off from extreme storms

Despite being short-lived, the changes we observed in bacterial community composition and environmental parameters during storms can nevertheless be detrimental to both the coastal ecosystem and human health. While most terrestrially-derived bacteria are benign to marine ecosystems, many are potentially pathogenic to corals and other marine organisms (Sutherland *et al*., 2004; Haapkylä *et al*., 2011; Pollock *et al*., 2014; Sheridan *et al*., 2014). Terrestrial run-off has been implicated in coral diseases, such as White Pox Disease and Red Band Disease, in many tropical and subtropical locations, including Madagascar (Haapkylä *et al*., 2011; Pollock *et al*., 2014; Sheridan *et al*., 2014), the Caribbean (Frias-Lopez *et al*., 2002; Patterson *et al*., 2002), and Australia’s Great Barrier Reef (Pollock *et al*., 2014). Furthermore, storm events can cause water-column mixing and sediment resuspension, leading to pathogenic marine bacteria that usually inhabit the seafloor to become more abundant in the water column (Hassard *et al*., 2016). Indeed, in our study we found several strains of *Vibrio spp*., *Pseudoalteromonas sp*., and Rhodobacteraceae bacteria specifically associated with coral disease (Sussman *et al*., 2008; Sheridan *et al*., 2014) to be significantly enriched in samples we collected during or following storm events, which could be caused by either terrestrial influence or marine resuspension (Supplemental Table 5). Considering the additional stress caused by turbidity and sedimentation during storm events, alongside potentially decreased pH and DO due to enhanced heterotrophic bacterial respiration (Weber *et al*., 2012; Altieri *et al*., 2017), corals and other organisms may be especially susceptible to pathogen infection during and after extreme storms.

Heavy rains and floods have long been implicated in transporting human pathogens (e.g. fecal coliform bacteria) to the marine environment (Pandey *et al*., 2014; De Jesus Crespo *et al*., 2019). Indeed, we found bacterial taxa that are potentially dangerous to humans—including Campylobacterales, Fusobacteriales, Bacillales, Clostridiales and Enterobacteriales (Bennett and Eley, 1993; Sharma *et al*., 2003; Davin-Regli *et al*., 2019)—significantly enriched during and following storms. Many of these bacteria were not detected in soil samples, suggesting additional sources of bacterial contamination in storm run-off. For instance, some of the Bacillales and Enterobacterales ASVs enriched in storm-influenced water samples were not present in soil samples and none of the enriched Clostridiales, Campylobacterales and Fusobacteriales ASVs were also found in soil samples (Figure 7). Likewise, these taxa were not found in mesocosms following soil addition in June or October, demonstrating the larger effect of storms and run-off on the coastal ecosystem than simply transporting sediment into the water. Human pathogens may derive from live-stock, storm drains, or overwhelmed waste treatment plants during storms (Weiskel *et al*., 1996; Jamieson *et al*., 2004). Interestingly, these taxa were already present in water samples collected on September 28—although they were more abundant on October 1 and 3—further indicating a cumulative effect of the typhoon season on the coastal ecosystem and emphasizing the context-dependency of storm effects. Ultimately, swimmers and other recreational users need to be aware that pathogenic bacteria are likely present in Okinawa coastal waters following large rain events.

## Conclusions & Future Directions

Despite challenges associated with sampling marine ecosystems during tropical storms and typhoons, this study describes the timing and nature of storm effects on coastal bacterial communities in Okinawa, Japan. We found that storm effects were transient, but highly context-dependent. The transient nature of storm effects should not be viewed as diminishing their potential impact on reef or human health. Differentially abundant bacteria during storms may cause disease in marine organisms and humans. We coupled controlled mesocosm studies with field observations in an effort to disentangle the effects of extreme wind and waves and enhanced currents from the effects of soil input during storms. While the mesocosm results were useful, future studies would benefit from more realistic run-off simulation than the soil additions we employed. Furthermore, it remains that we did not perform bacterial cell counts or otherwise measure bacterial biomass or metabolic activity, thus leaving the possibility that run-off increased microbial biomass or differentially affected microbial physiology. Future studies may capture such responses by measuring bacterial respiration rates or by performing metatranscriptomics.

It is important to note that environmental effects of extreme storms will vary in terms of intensity, spatial extent and duration in different ecosystems and need to be evaluated locally (Paerl *et al*., 2006; Zhang *et al*., 2009; Herbeck *et al*., 2011). Storm effects were transient in the open, tidally-flushed Okinawa coast, but more prolonged storm effects have been observed in other coastal systems, particularly semi-enclosed areas, such as bays and estuaries, where terrestrial sediment loads can have residence times from weeks to years (Paerl *et al*., 2001; Zhang *et al*., 2009; Herbeck *et al*., 2011). Therefore, we suggest that the short-term study of typhoon events follow an adaptive sampling strategy (Wetz and Paerl, 2008), which involves the definition of well-established baselines for various physicochemical and biological parameters. Regional monitoring programs, including a comprehensive understanding of background conditions, are essential for a better interpretation of ecological consequences of extreme storm events.

## Experimental Procedures

### Study setting

Seawater was collected for metabarcoding and physicochemical analysis from four nearshore sampling points approximately 250–500 m apart, along the central west coast of Okinawa Island—a semi-urban region with mixed land-use, including agriculture and coastal development projects (Supplemental Figure 2). The sampling points were each at least 1.2 km from the nearest concentrated fresh-water input (e.g. streams or rivers). At the start of the 2018 typhoon season, samples were collected before (June 13), during (June 16), and after (June 19) tropical storm Gaemi, which struck Okinawa on June 16, 2018 and caused a red soil pollution event (Figure 1, Supplemental Figure 3A). Towards the end of the typhoon season, samples were collected before (Sept 28), during (Oct 1 and Oct 3) and after (Oct 8) a red soil pollution event caused by typhoon Trami, which made landfall with Okinawa on September 30, and was prolonged by Typhoon Kong-Rey, which approached Okinawa on October 5th (Figure 1, Supplemental Figure 3B–C).

### Seawater sampling for DNA and physicochemical analysis

Surface seawater was collected for DNA metabarcoding by submerging clean 500 mL Nalgene bottles just below the sea surface. Seawater for dissolved Fe (dFe) and nutrient analysis was collected in acid-cleaned 50 mL Falcon tubes. Physicochemical properties—dissolved oxygen (DO), salinity, temperature, and turbidity—were measured with a CTD RINKO profiler (JFE Advantech, Japan) at each site. After being immediately transported to the laboratory, seawater samples for metabarcoding were filtered through 0.2 µm pore-size Polytetrafluoroethylene (PTFE) filters (Millipore) under gentle vacuum and filters were stored at −20 °C for later DNA extraction. Seawater samples for dFe and nutrient analysis were filtered through 0.45 µm pore-size acid-washed Teflon digiFILTERS (SPC Science, Canada) and filtered water samples were stored at −20 °C for later chemical analysis.

### Mesocosm experimental design

Seawater was collected for concurrent mesocosm experiments on June 11 (26.512 °N, 127.872°E) and October 10 (26.479 °N, 127.829 °E). Nearshore coastal seawater was pumped from just below the sea surface and filtered through 1 mm and 300 µm nylon mesh sizes to remove debris and larger organisms. Acid-cleaned 22 L clear-plastic carboys were rinsed twice with seawater before filling to 20 L with filtered seawater. Bottles were covered with parafilm and kept shaded during transport to the Okinawa Institute of Science and Technology (OIST) Marine Science Station, where they were submerged in a basin with continuous flow-through seawater to keep conditions within the bottles similar to natural conditions. Mesocosm bottles were topped with silicone sponge stoppers to allow gas exchange, but limit evaporation and prevent dust, water or other contaminants from entering the bottles during the experiment (Supplemental Figure 4). In addition, small pumps were included in each mesocosm to maintain water circulation (2 L min^-1^). HOBO temperature and light loggers (Onset) were fastened to the pumps and at the same depth in the basin surrounding mesocosms to ensure that mesocosms conditions remained similar to ambient conditions (Supplemental Figure 5). In addition, salinity and DO were measured each time water was sampled from the mesocosms throughout the experiment (Supplemental Figure 6). Mesocosms experiments included 9–10 mesocosms: 4–5 control replicates (four in June and five in October) and 5 treatments replicates with red soil added to an ecologically relevant concentration of 200 mg L^-1^ (O’Connor *et al*., 2016). Mesocosms were sampled (100 mL for metabarcoding, 50 mL for dFe and nutrient concentration) with 50 mL sterile pipettes before the experiment started (t0), and 1, 4, 12, 24, and 48 h following red soil addition to treatment bottles. Water samples were processed as described in the previous section.

### Red soil collection for mesocosm experiments and DNA analysis

Soil samples were collected from an open agricultural field with exposed soil (26.507 °N, 127.868 °E), on June 10 and October 9 for addition to red soil treated mesocosms and to evaluate soil microbiomes. Soil samples were sieved through 330 μm mesh and maintained at 4 °C for 24 h until use in the mesocosm experiment. To determine soil moisture content, 10 g subsamples (n=10) were weighed and dried at 100 °C by following the standard method AS 1289.2.1.1-2005 (Standards Association of Australia). Soil moisture content was used to calculate how much wet soil should be added to mesocosms in order to reach the final concentration of 200 g soil (dry weight) per L seawater. In both June (n=4) and October (n=2), additional 50 g aliquots of soil were kept at −20 °C for subsequent metabarcoding analysis.

### Chemical analysis: dFe and major nutrients

Dissolved Fe (dFe) concentration was determined following the methodology of Wu and Boyle (1998). This method uses a Mg (OH)_2_ co-precipitation to pre-concentrate Fe from seawater followed by an isotope dilution method. Seawater dFe was quantified using an internal standard element (^57^Fe) with inductively coupled plasma mass spectrometry (Element 2, Thermo Scientific). The mass spectrometer was operated in medium-resolution mode with 4000 resolution (FWHM). The mass calibration was performed using a multi-element ICP-MS tune-up solution (Thermo Fisher Scientific). In order to ensure the quality of the ICP-MS analysis, control standards and samples (i.e. analytical replicates, certified reference material and analytical blank) were analyzed once every 12 samples. Recovery of Fe from reference certified material QC3163 (Sigma-Aldrich, USA) was satisfactory and ranged from 65 to 80%. The overall error associated with the analytical process was typically lower than 5% and never higher than 15%. All measurements were above the instrument’s detection limit. Analysis was carried out at the OIST Instrumental Analysis Section mass spectrometry laboratory. Special attention was paid to avoid Fe contamination and an exhaustive cleaning process was carried out following the methods of (King and Barbeau, 2011).

Nutrient concentrations—including Nitrate (NO_3_^-^), Nitrite (NO_2_^-^), Ammonium (NH^4+^), Phosphate (PO_43_^-^) and Silica (SiO_2_)—were determined on a QuAAtro39 Continuous Segmented Flow Analyzer (SEAL Analytical) following manufacturer guidelines. Final concentrations were calculated through AACE software (SEAL Analytical). Nutrient Analysis was carried out at the Okinawa Prefecture Fisheries and Ocean Technology Center.

### Nutrient and dFe statistical analyses

Mesocosm data were found to be normally distributed with a Shapiro-Wilks test and, therefore, a one-way ANOVA was performed to test overall differences between treatments. Post-hoc Tukey HSD analysis was performed to identify which specific groups differed. Field data were not normally distributed, regardless of transformation, so a Kruskal-Wallis test was used to test for significant differences between sampling dates. Analyses were performed within the R statistical environment (R Core Team 2018).

### DNA extraction and metabarcode sequencing

DNA was extracted from frozen PTFE filters following the manufacturer protocol for the DNeasy PowerWater Kit (Qiagen), including the optional heating step. DNA was extracted from soil samples by following manufacturer protocol for the DNeasy PowerSoil Kit (Qiagen). Metabarcode sequencing libraries were prepared for the V3/V4 region of the bacterial 16S ribosomal RNA gene following Illumina’s “16S Metagenomic Sequencing Library Preparation” manual without any modifications. Sequencing libraries were transferred to the OIST Sequencing Center for 2×300-bp sequencing on the Illumina MiSeq platform with v3 chemistry. Overall, 18.4 million sequencing reads were generated in this study, with 76,217–219,584 sequencing reads per sample (mean = 137,585). Sequencing data are available from the NCBI Sequencing Read Archive under the accession PRJNA564579.

### Metabarcode analyses

Sequencing reads were denoised using the Divisive Amplicon Denoising Algorithm (Callahan *et al*., 2016) with the DADA2 plug-in for QIIME 2 (Bolyen *et al*., 2019). Following denoising, 11.8 million sequences remained with 11,061–153,646 sequences per sample (mean = 88,033). A total of 36,007 ASVs were identified in our dataset, with 64–2,886 unique ASVs observed per sample (mean = 642). Taxonomy was assigned to representative ASVs using a Naive Bayes classifier trained on the SILVA 97% consensus taxonomy (version 132, (Quast *et al*., 2013) with the QIIME 2 feature-classifier plug-in (Bokulich *et al*., 2018). The results were imported into the R statistical environment (R Core Team 2018) for further analysis with the Bioconductor phyloseq package (McMurdie and Holmes, 2013). The ASV richness for each sample was estimated using the R package breakaway (Willis and Bunge, 2015) and differences in estimated richness between sample types were tested for with the betta function (Willis *et al*., 2017). In order to minimize compositional bias inherent in metabarcoding data as much as possible, we used the Aitchison distance between samples (Gloor *et al*., 2017) for principal coordinate analyses (PCoA). Permutational analyses of variance (PERMANOVA) on Aitchison distances were performed with the adonis function (999 permutations) in the R package vegan (Oksanen *et al*., 2019) to test whether shifts in community composition were statistically significant. Lastly, we used the DESeq2 bioconductor package (Love *et al*., 2014) to determine which ASVs were significantly differentially abundant (False Discovery Rate adjusted *p*-value < 0.05) in water samples collected from field sites before, during, and after storms. We then checked if significantly differentially abundant ASVs were also present in soil samples from respective months to assess whether differentially abundant ASVs were soil-derived. Intermediate data files and the code necessary to replicate analyses are available in a GitHub repository: https://github.com/maggimars/RedSoil.

## Acknowledgments

We thank Akinori Murata and Marine Le Gal for help with sample collection, Kazumi Inoha, Koichi Toda, Kosuke Mori and Okinawa Marine Science Support Section, OIST for assistance with experimental design and technical support and Koichi Kinjo from the Okinawa Prefectural Institute of Health and Environment for his advice. We also thank OIST sequencing center for DNA sequence support and Okinawa Prefecture Fisheries and Ocean Technology Center for nutrient analysis assistance. MMB was supported by a Japan Society for the Promotion of Science (JSPS) DC1 graduate student fellowship. This work was supported by the JSPS KAKENHI program [Early-Career Project No. 18K18203] and the OIST Marine Biophysics Unit. Furthermore, we confirm that we have no conflicts of interest to disclose.

