## Supplemental material for "Extreme storms cause rapid, context-dependent shifts in nearshore subtropical bacterial communities"

**List of supplemental files**

Supplemental Figure 1. Photographs of red soil run-off in nearshore waters of the central west coast of Okinawa, Japan

Supplemental Figure 2. Map of sampling sites in Okinawa

Supplemental Figure 3. Tracks of tropical cyclones monitored in this study

Supplemental Figure 4. Mesocosm experimental design

Supplemental Figure 5. Light intensity and temperature in mesocosms

Supplemental Figure 6. Dissolved oxygen and salinity in mesocosms

Supplemental Figure 7. Temporal variation of micro- and macro-nutrient concentrations in a mesocosm experiment following red soil addition in October, 2018

Supplemental Figure 8. Principal coordinates analysis (PCoA) of Aitchison distances between bacterioplankton community composition before, during, and after storm events in June and October, 2018.

Supplemental Figure 9. Richness estimates for bacterial communities in control and red soil amended bottles in mesocosm experiments

Supplemental Figure 10. Principal coordinates analysis (PCoA) of Aitchison distances between bacterial community compositions in mesocosms

Supplemental Table 1. Micro- and macro-nutrient concentrations before, during, and after storm events in June and October, 2018

Supplemental Table 2. Micro- and macro-nutrient concentrations in a mesocosm experiment following red soil addition in October, 2018

Supplemental Table 3. Results from Kruskall-Wallis hypothesis testing for nutrient concentrations from field samples

Supplemental Table 4. Results from two-way ANOVA hypothesis testing for nutrient concentrations in mesocosms

Supplemental Table 5. Log2 fold change for Amplicon Sequence Variants (ASVs) significantly enriched in storm samples with significant blast hits to known coral pathogens

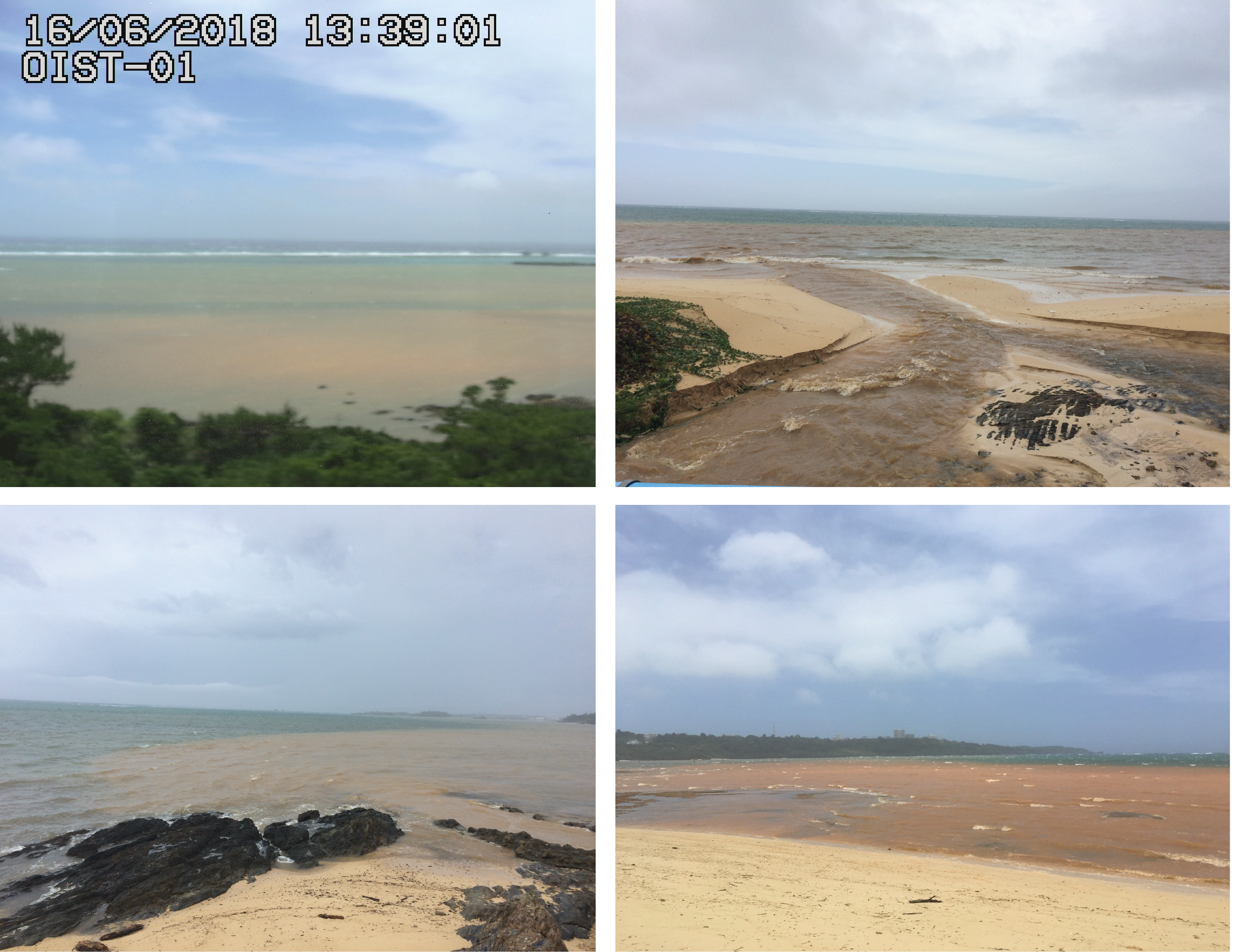

**Supplemental Figure 1.** Photographs showing the red soil run-off after the tropical storm Gaemi the 16th of June of 2018. Photographs were taken within a 10 km radius of the study area between 13:00 and 15:30 pm, just 2–3 hours after the peak of rain.

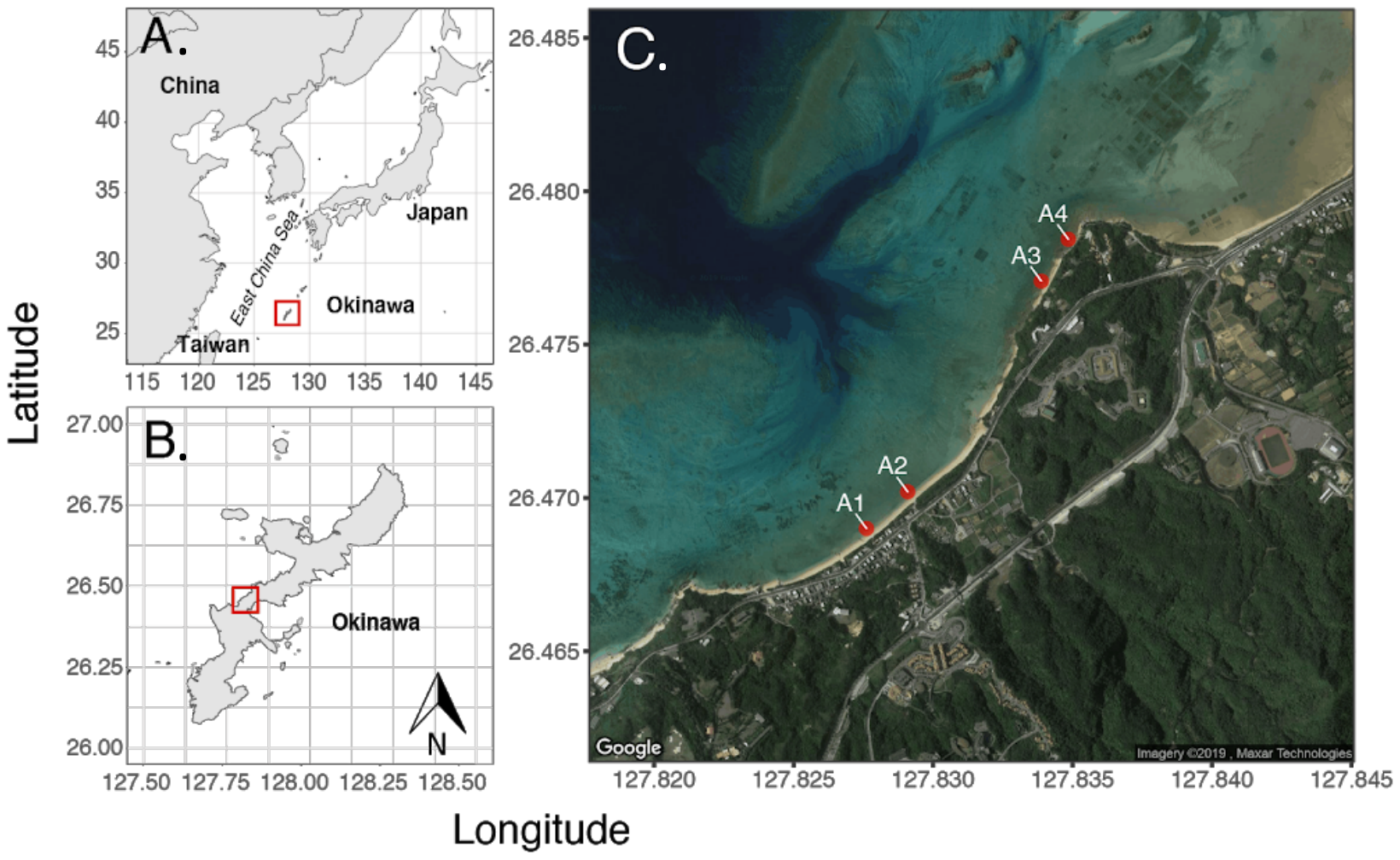

**Supplemental Figure 2.** A) Map showing the location of Okinawa Island in the East China Sea. B) Map showing the location of the central-west coast of Okinawa Island. C) Location of the 4 nearshore sampling points in Onna-son, on the central-west coast of Okinawa Island.

A. Tropical Storm Gaemi (June 16, 2018)

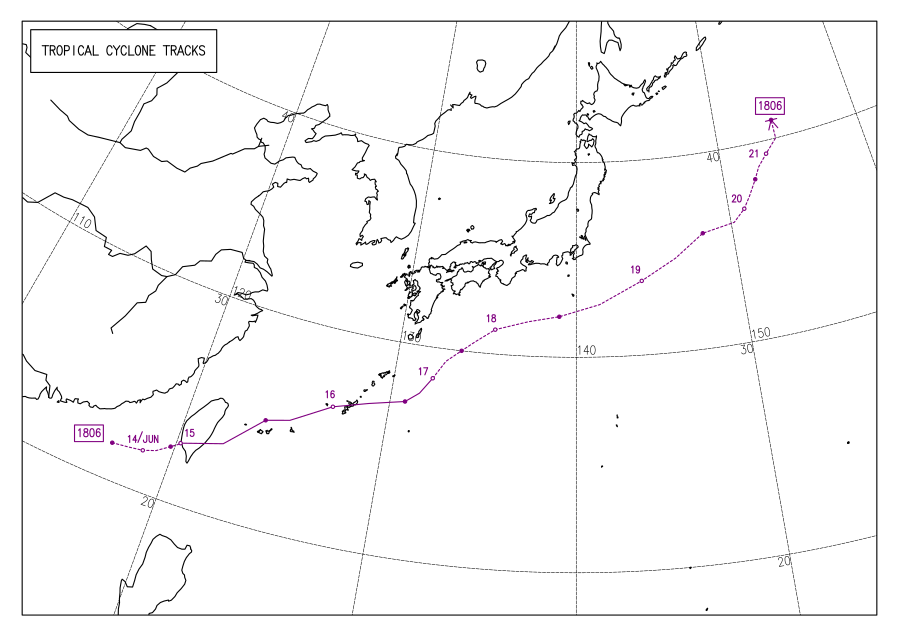

B. Typhoon Trami (September 29–30, 2018)

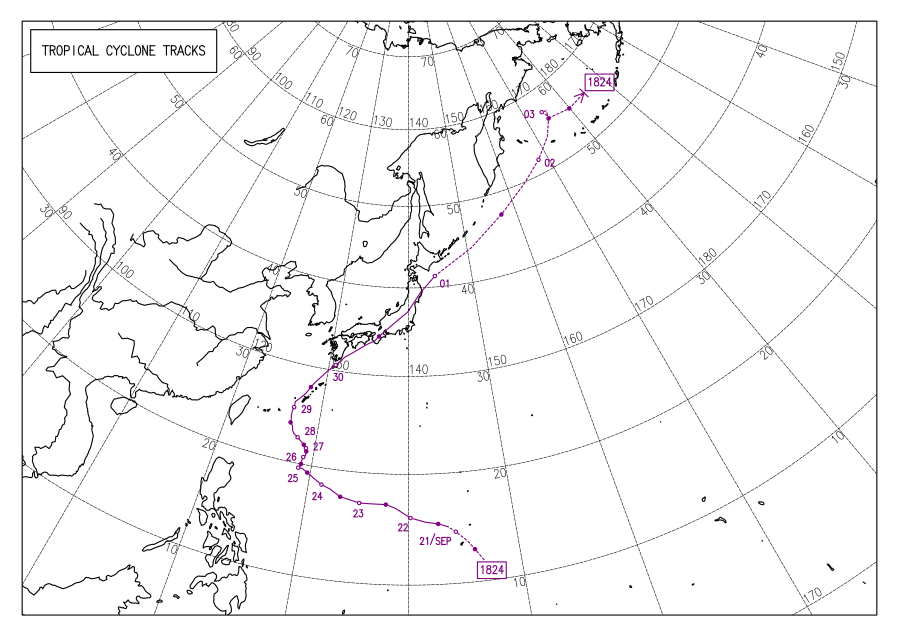

C. Typhoon Kong-Rey (October 4–5, 2018)

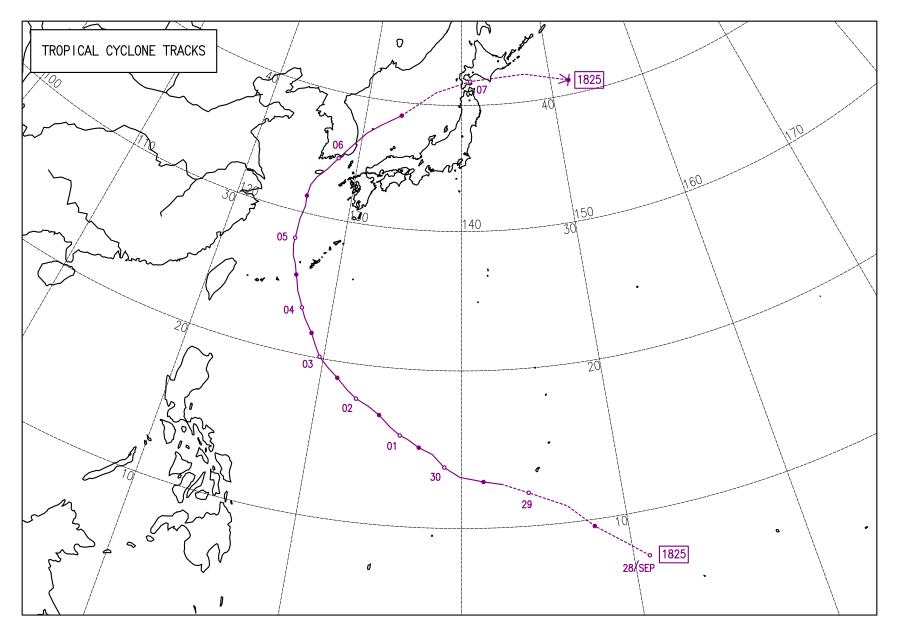

**Supplemental Figure 3.** Tropical Cyclone track information for the two storm events monitored during this study: tropical storm Gaemi, which made landfall with Okinawa Island on June 16 (A), two successive typhoons, Trami (B) and Kong-Rey (C), which made landfall on September 30 and October 5, 2018, respectively. Maps were obtained from the Japan Meteorological Agency (JMA, Tropical Cyclone Analysis Archive).

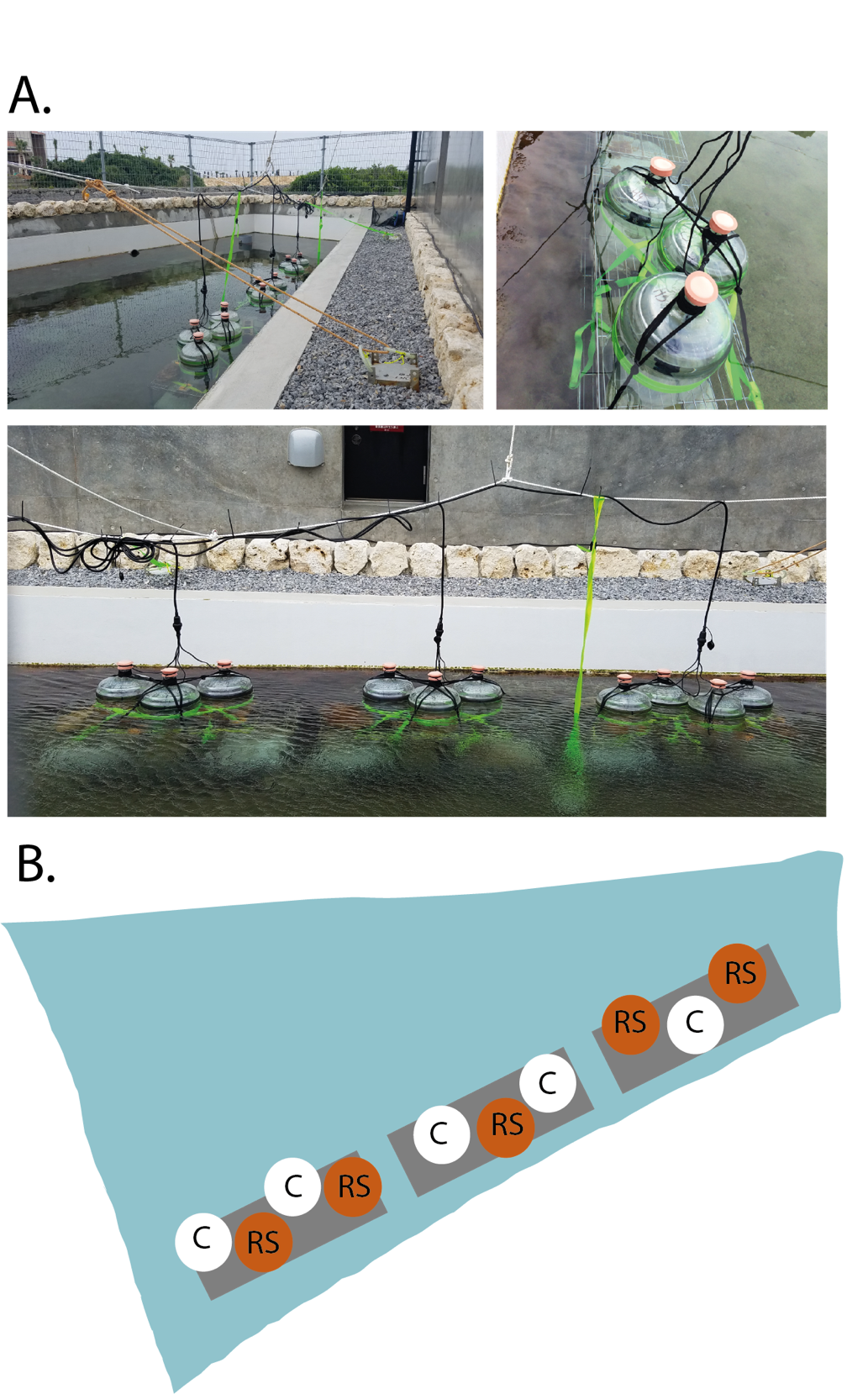

**Supplemental Figure 4.** A) photographs and B) sketch of the mesocosm experimental set-up at the OIST Seragaki Marine Station. Mesocosms were used to determine the time course impact of red soil pollution on biological communities and physicochemical properties of the water.

**Supplemental Figure 5. Mean temperature (°C) and light intensity (Lux) during the mesocosms experiments in June (A) and October (B) during daylight and nighttime hours.** Temperature and Light intensity were monitored inside bottles and outside the bottles in the surrounding basin to ensure that mesocosms conditions remained similar to ambient conditions. Data were collected every 30 min. throughout the experiment by HOBO temperature and light loggers (Onset). Error bars represent one standard deviation. (Day hours = 04:30-19:00; Night hours: 19:00-04:30).

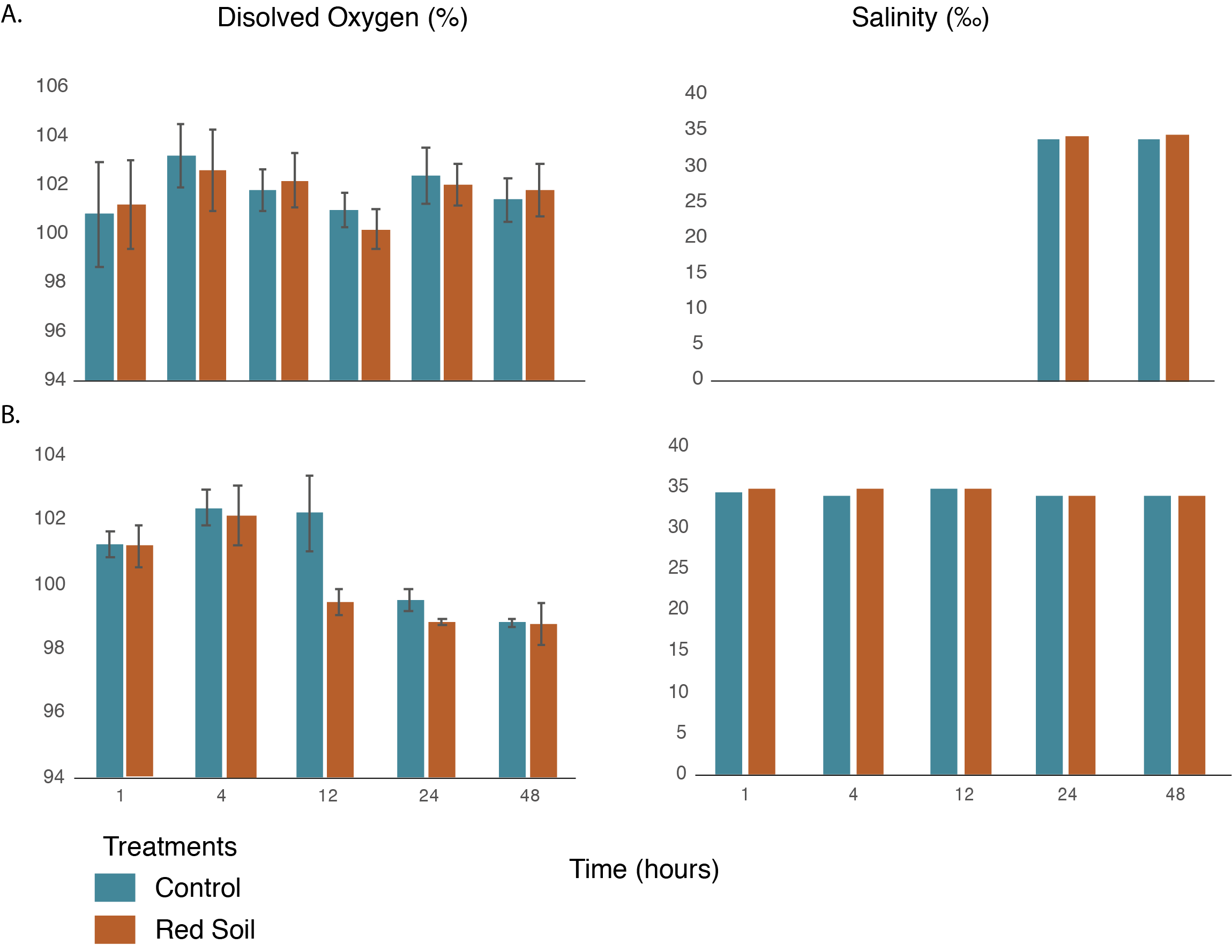

**Supplemental Figure 6. Mean Dissolved Oxygen (in %) and salinity (‰) in mesocosm experiments in June (A) and October (B).** Dissolved oxygen (DO) and salinity were monitored inside the mesocosm bottles at each sampling time-point: 1, 4, 12, 24, and 48 hours after the start of the experiment. DO was measured using a Portable optical DO meter (ARO-PR, JFE Advantech Co., Ltd) and salinity was assessed with a [Master-500 Refractometer](https://www.thomassci.com/search/go?p=R&ts=custom&uid=537074483&w=Refractometer%20Atago&method=and&sid=2&isort=score&url=http%3a%2f%2fwww.thomassci.com%2fInstruments%2fRefractometers%2f_%2fMaster-500-Refractometer&rsc=XfOq-r9XgI9saEib&domainSpecificUrl=https%3a%2f%2fwww.thomassci.com%2fInstruments%2fRefractometers%2f_%2fMaster-500-Refractometer) (Atago). Error bars represent one standard deviation.

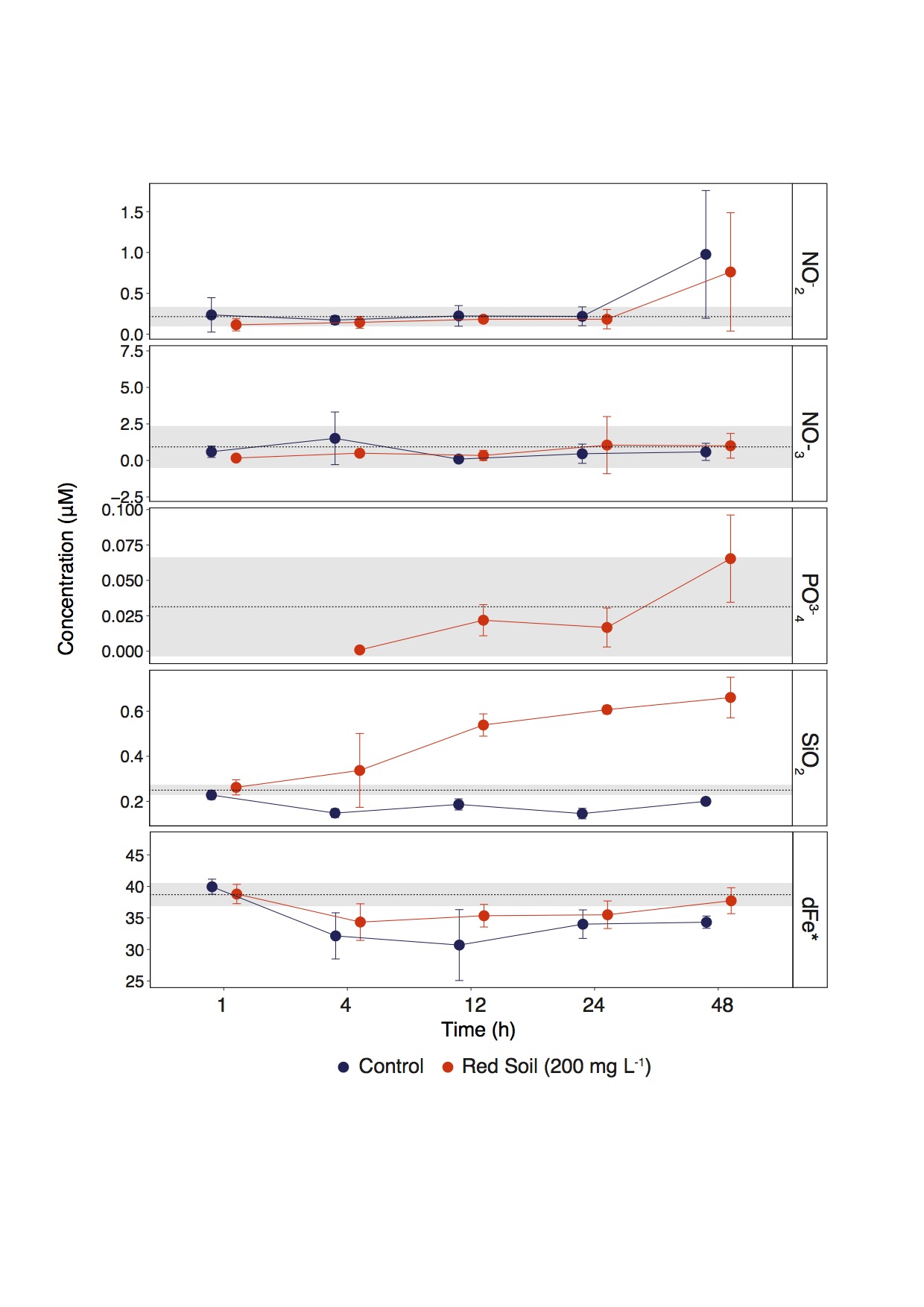

**Supplemental Figure 7. Temporal variation of micro- and macro-nutrient concentrations in mesocosm experiment performed in October, 2018.** Points represent mean concentrations of dissolved NO_2_^-^, NO_3_^-^, PO_4_^3-^, SiO_2_ and dFe (* dFe concentration is in nM) in control (blue points) and red soil treated (red points) mesocosms. Samples for nutrient analysis were taken at t0 and 1, 4, 12, 24, and 28 h after red soil (200 mg/L) was added to treatment mesocosms. Error bars on points denote ± one standard deviation. The dashed black horizontal line is the mean value for all mesocosms at t0 and the shaded region represents ± one standard deviation. Only values higher than the limit of detection are shown in the figure; PO_4_^3-^ concentrations stayed below the limit of detection in control mesocosms after t0.

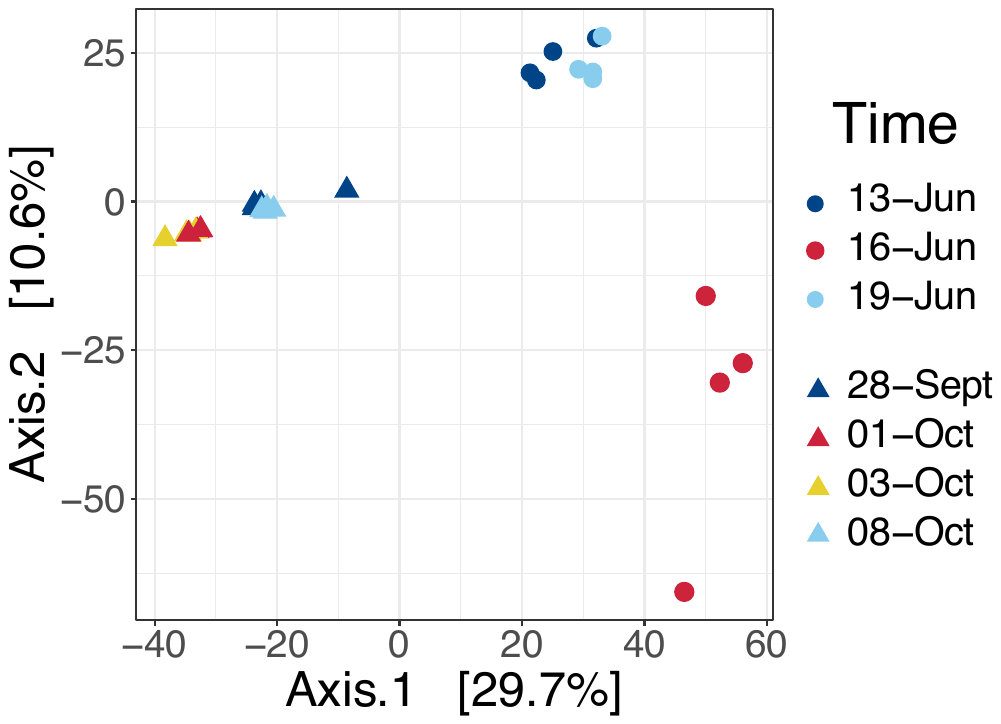

**Supplemental Figure 8. Principal coordinates analysis (PCoA) of Aitchison distances between bacterioplankton community composition before, during, and after storm events in June and October, 2018.** 13-Jun was before the June storm, 16-Jun was during, and 19-Jun was after. For the October event, 28-Sept. was before the storms, 01-Oct. and 03-Oct. were between, and 08-Oct. was after. Points are colored by sampling date, with cool colors representing dates before and after storms and warm colors representing dates during or between storms. Point shape represents sampling month, with June samples represented by circled and October samples represented by triangles. Samples clustered most strongly by sampling month. PERMANOVA results when performed by sampling month were statistically significant (*p* < 0.01, *F* = 5.37, *R*^2^ = 0.17).

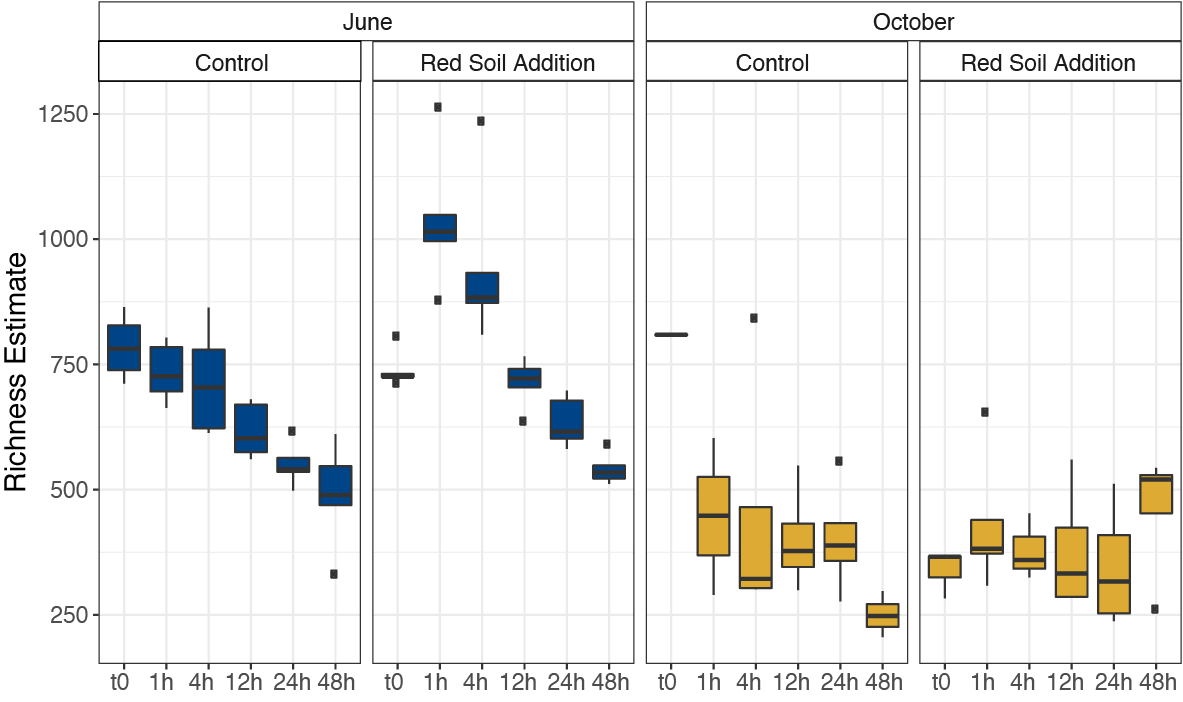

**Supplemental Figure 9. Richness estimates for bacterial communities in control and red soil amended bottles in mesocosm experiments conducted in June and October, 2018.** ASV richness in red soil treated and control mesocosms was estimated using the R package Breakaway. Twenty-Liter mesocosms were set up in ambient conditions with natural seawater collected near Okinawa in June and October 2018 and sampled at t0, t1–t5: 1, 4, 12, 24, and 48 hours after initiating experimental conditions. ASV richness was higher at the start of the June experiment than at the start of the October experiment and red soil addition caused an increase in ASV richness in the June but not in the October.

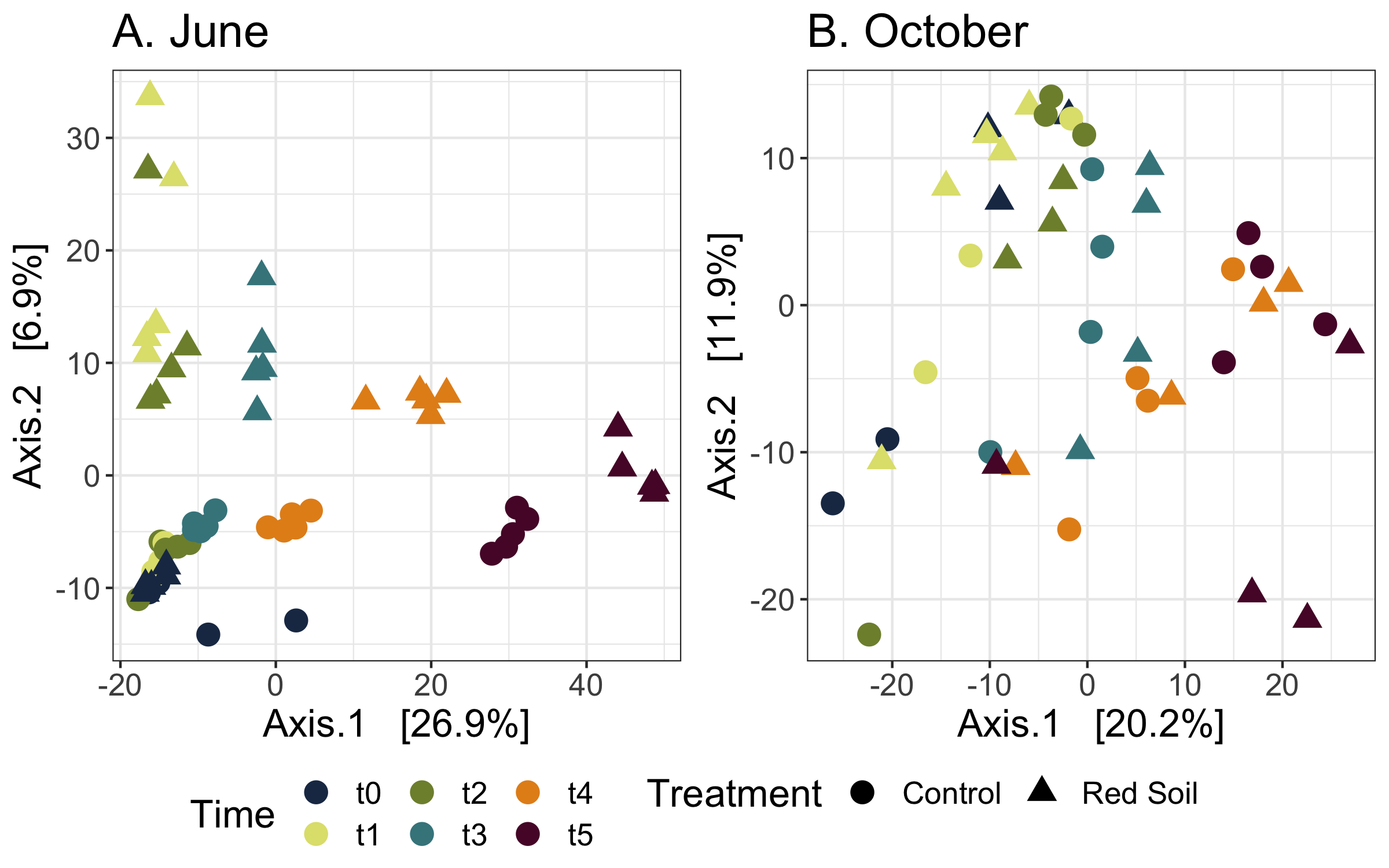

**Supplemental Figure 10. Principal coordinates analysis (PCoA) of Aitchison distances between bacterioplankton community composition in mesocosm experiments conducted in June (A) and October (B), 2018.** Twenty-Liter mesocosms were set up in ambient conditions with natural seawater collected near Okinawa in June and October 2018 and sampled at t0, before red soil addition and t1–t5: 1, 4, 12, 24, and 48 hours after red soil addition. Point color indicates sampling time and point shape corresponds to treatment: circles represent control bottles and triangles represent red soil treated bottles, to which 200 mg L^-1^ freshly collected Red Soil were added. In June (A), red soil addition caused t1 samples to diverge from t0 samples, with the distance between treatments decreasing over time. Conversely, red soil addition in October (B) did not cause a clear shift in the bacterial community within mesocosms. Samples clustered along the primary axis by sampling time in both treatments during both experiments, although this effect was stronger for the June experiment.

**Supplemental Table 1.** Micro- and macro-nutrient field concentrations before, during, and after storm events in June and October, 2018

| **Date** | **month** | **NO_2_^-^ μM** | **NO_3_^-^ μM** | **NH_4_^+^ μM** | **PO_4_^3-^ μM** | **SiO_2_ μM** | **dFe nM** |
| --- | --- | --- | --- | --- | --- | --- | --- |
| **13/06/2019** | June | 0.175 | 0.515 | 5.770 | 0.492 | 0.290 | 37.535 |
| **13/06/2019** | June | 0.122 | 12.798 | 1.618 | 0.226 | 2.904 | 38.807 |
| **13/06/2019** | June | 0.227 | 1.308 | 0.961 | 0.076 | 0.934 | 36.648 |
| **13/06/2019** | June |  |  |  |  |  | 37.939 |
| **14/06/2019** | June | 0.300 | 0.186 | 0.479 | 0.025 | 4.148 |  |
| **14/06/2019** | June | 0.434 | 1.965 | 1.457 | 0.054 | 3.526 |  |
| **14/06/2019** | June | 0.298 | 0.168 | 0.427 | 0.037 | 6.484 |  |
| **14/06/2019** | June | 0.179 | 0.085 | 0.643 | 0.019 | 2.205 |  |
| **15/06/2019** | June | 0.243 | 0.344 | 0.392 | 0.028 | 10.773 |  |
| **15/06/2019** | June | 0.330 | 0.326 | 0.082 | 0.024 | 12.732 |  |
| **15/06/2019** | June | 0.554 | 0.456 | 0.996 | 0.021 | 2.173 |  |
| **15/06/2019** | June | 0.523 | 0.404 | 0.656 | 0.028 | 5.322 |  |
| **16/06/2019** | June | 0.497 | 0.796 | 1.054 | 0.086 | 25.435 | 37.742 |
| **16/06/2019** | June | 0.163 | 0.572 | 0.799 | 0.066 | 15.974 | 39.763 |
| **16/06/2019** | June | 0.476 | 11.570 | 4.302 | 0.539 | 13.240 | 66.128 |
| **16/06/2019** | June | 0.449 | 0.999 | 1.292 | 0.159 | 10.077 | 37.661 |
| **18/06/2019** | June | 0.649 | 0.578 | 16.541 | 0.346 | 3.372 | 38.010 |
| **18/06/2019** | June | 0.132 | 0.978 | 2.033 | 0.102 | 6.748 | 45.812 |
| **18/06/2019** | June | 0.320 | 0.086 | 0.340 | 0.039 | 0.436 | 39.188 |
| **18/06/2019** | June | 0.469 | 0.091 | 0.288 | 0.018 | 1.171 | 36.234 |
| **26/06/2019** | June | 0.002 | 0.048 | 0.336 | 0.003 | 2.680 |  |
| **26/06/2019** | June | 0.014 | 0.469 | 0.694 | 0.013 | 17.817 |  |
| **26/06/2019** | June |  |  | 0.813 | 0.011 |  |  |
| **26/06/2019** | June | 0.004 |  | 1.161 | 0.033 |  |  |
| **28/09/2019** | October | 0.087 | 1.188 | 0.516 | 0.038 | 12.287 | 67.161 |
| **28/09/2019** | October | 0.050 | 0.837 | 0.448 | 0.017 | 8.863 | 38.569 |
| **28/09/2019** | October | 0.021 | 1.105 | 0.351 | 0.021 | 2.138 | 34.058 |
| **28/09/2019** | October | 0.056 | 2.493 | 0.260 | 0.008 | 0.820 | 35.668 |
| **01/10/2019** | October | 0.073 | 1.806 | 0.541 | 0.026 | 10.020 | 33.691 |
| **01/10/2019** | October | 0.085 | 1.055 | 0.400 | 0.020 | 13.692 | 39.391 |
| **01/10/2019** | October | 0.037 | 2.676 | 0.218 | 0.007 |  | 34.978 |
| **01/10/2019** | October | 0.087 | 1.048 | 1.675 | 0.006 | 2.192 | 36.045 |
| **03/10/2019** | October | 0.194 | 1.063 | 0.110 | 0.002 | 1.316 | 31.412 |
| **03/10/2019** | October | 0.341 | 0.542 | 0.263 | 0.003 | 6.190 | 35.232 |
| **03/10/2019** | October | 0.191 | 0.667 | 0.116 |  | 1.054 | 43.491 |
| **03/10/2019** | October | 0.404 | 0.515 | 0.498 |  | 3.271 | 36.160 |
| **07/10/2019** | October | 0.254 | 1.092 |  |  | 9.909 | 36.905 |
| **07/10/2019** | October | 0.287 | 0.482 | 0.013 | 0.008 | 64.089 | 37.563 |
| **07/10/2019** | October | 0.193 | 0.380 | 1.155 | 0.005 | 13.332 | 36.288 |
| **07/10/2019** | October | 0.241 | 0.129 | 3.337 | 0.006 | 2.804 | 36.032 |

**Supplemental Table 2.** Micro- and macro-nutrient concentrations in mesocosm experiment performed in October, 2018. Sample IDs correspond to sampling time point and treatment; C for control and R for red soil addition.

| **ID** | **NO_3_^-^ + NO_2_^-^**  **(μM)** | **NO_2_^-^ (μM)** | **NO_3_^-^ (μM)** | **PO_4_^3-^ (μM)** | **SiO_2_ (μM)** | **dFe (nM)** |
| --- | --- | --- | --- | --- | --- | --- |
| **t0** | 0.191 | 0.135 | 0.126 | 0.071 | 0.233 | 41.308 |
| **t0** | 0.258 | 0.106 | 0.349 | 0.021 | 0.238 | 38.870 |
| **t0** | 1.822 | 0.374 | 3.096 | 0.003 |  | 36.315 |
| **t0** | 0.242 | 0.245 | 0.144 |  | 0.279 | 37.058 |
| **t1_C** | 0.461 | 0.155 | 0.622 |  | 0.245 | 38.247 |
| **t1_C** | 0.676 | 0.116 | 1.062 |  | 0.246 | 40.393 |
| **t1_C** | 0.108 | 0.551 | 0.103 |  | 0.209 | 41.033 |
| **t1_C** | 0.582 | 0.122 | 0.605 |  | 0.213 | 40.115 |
| **t1_R** | 0.149 | 0.127 | 0.173 |  | 0.294 | 37.767 |
| **t1_R** | 0.131 | 0.202 | 0.134 |  | 0.257 | 38.682 |
| **t1_R** | 0.204 | 0.107 | 0.206 |  | 0.280 | 40.995 |
| **t1_R** | 0.134 | 0.021 | 0.155 |  | 0.218 | 37.729 |
| **t2_C** | 0.103 | 0.122 | 0.164 |  |  | 37.445 |
| **t2_C** | 0.879 | 0.245 | 1.440 |  |  | 31.508 |
| **t2_C** | 2.446 | 0.175 | 4.080 |  | 0.134 | 29.117 |
| **t2_C** | 0.285 | 0.148 | 0.366 |  | 0.162 | 30.551 |
| **t2_R** | 0.210 | 0.113 | 0.234 |  | 0.444 | 37.630 |
| **t2_R** | 0.467 | 0.212 | 0.730 |  | 0.095 | 32.131 |
| **t2_R** | 0.427 | 0.197 | 0.587 | 0.001 | 0.427 | 33.261 |
| **t2_R** | 0.337 | 0.059 | 0.441 |  | 0.384 |  |
| **t3_C** | 0.032 | 0.222 | 0.013 |  | 0.179 | 24.865 |
| **t3_C** | 0.128 | 0.395 | 0.057 |  | 0.167 | 36.077 |
| **t3_C** | 0.176 | 0.183 | 0.013 |  |  | 31.153 |
| **t3_C** | 0.238 | 0.094 | 0.278 |  | 0.214 |  |
| **t3_R** | 0.510 | 0.138 | 0.670 | 0.009 | 0.593 | 34.329 |
| **t3_R** | 0.069 | 0.205 | 0.019 | 0.019 | 0.538 | 34.947 |
| **t3_R** | 0.441 | 0.224 | 0.616 | 0.024 | 0.550 | 34.145 |
| **t3_R** | 0.125 | 0.165 | 0.052 | 0.035 | 0.474 | 37.994 |
| **t4_C** | 0.082 | 0.154 | 0.020 |  | 0.130 | 32.128 |
| **t4_C** | 0.082 | 0.391 | 0.028 |  | 0.123 | 33.978 |
| **t4_C** | 0.979 | 0.178 | 1.415 |  | 0.161 | 37.204 |
| **t4_C** | 0.291 | 0.150 | 0.375 |  | 0.170 | 32.746 |
| **t4_R** | 0.062 | 0.138 | 0.008 | 0.009 | 0.611 | 37.963 |
| **t4_R** | 0.160 | 0.053 | 0.176 | 0.024 | 0.587 | 34.501 |
| **t4_R** | 2.307 | 0.210 | 3.979 | 0.002 |  | 36.529 |
| **t4_R** | 0.104 | 0.332 | 0.025 | 0.032 | 0.622 | 33.015 |
| **t5_C** | 0.207 | 1.882 | 0.114 |  | 0.209 | 33.153 |
| **t5_C** | 1.487 | 0.410 | 1.195 |  | 0.186 | 33.932 |
| **t5_C** | 0.208 | 1.373 | 0.058 |  | 0.195 | 34.908 |
| **t5_C** | 1.149 | 0.243 | 0.983 |  | 0.211 | 35.314 |
| **t5_R** | 0.585 | 0.341 | 0.941 | 0.029 | 0.717 | 37.419 |
| **t5_R** | 0.242 | 0.036 | 0.168 | 0.051 | 0.688 | 40.702 |
| **t5_R** | 0.433 | 1.662 | 0.727 | 0.083 | 0.527 | 36.670 |
| **t5_R** | 1.956 | 1.012 | 2.174 | 0.097 | 0.710 | 36.121 |

**Supplemental Table 3**. Results of non-parametric Kruskal-Wallis tests comparing the concentrations of dissolved nutrients in the field (NO_2_^-^, NO_3_^-^, NH_4_^+^, PO_4_^3-^, SiO_2_, dFe) before, during, and after storm events in June and October, 2018. Significant *p*-values (*p* < 0.05) are marked with a * in the significance column and nonsignificant values are marked n.s.

| **Source** | **Variable** | **Chi-squared** | | **N** | ***p*** | **Sig** |
| --- | --- | --- | --- | --- | --- | --- |
| June | NO_2_^-^ | 11.943 | 5 | | 0.0356 | * |
|  | NO_3_^-^ | 8.8701 | 5 | | 0.144 | n.s. |
|  | NH_4_^+^ | 6.413 | 5 | | 0.2681 | n.s. |
|  | PO_4_^3-^ | 11.583 | 5 | | 0.0123 | * |
|  | SiO_2_ | 11.403 | 5 | | 0.0439 | * |
|  | dFe | 1.5 | 2 | | 0.4724 | n.s. |
| October | NO_2_^-^ | 11.515 | 3 | | 0.00924 | * |
|  | NO_3_^-^ | 7.8088 | 3 | | 0.05013 | n.s. |
|  | NH_4_^+^ | 2.3792 | 3 | | 0.4975 | n.s. |
|  | PO_4_^3-^ | 7.8297 | 3 | | 0.0496 | * |
|  | SiO_2_ | 4.65 | 3 | | 0.1993 | n.s. |
|  | dFe | 1.1471 | 3 | | 0.7657 | n.s. |

**Supplemental Table 4.** Results of the two-way ANOVA tests comparing dissolved nutrient concentrations (NO_2_^-^, NO_3_^-^, PO_4_^3-^, SiO_2_, dFe) in control and red soil treated mesocosms during the October experiment. DF= degrees of freedom; SS = sum of squares; *p* = *p*-value and Sig = significance. Significant *p-*values (*p* < 0.05) are shown as follows in the significant column: ‘***’ = 0.001; ‘**’ = 0.01; ‘*’ = 0.05; ‘n.s.’ = not significant.

| **Source of Variation** | | **Dependent Variable** | **DF** | **SS** | ***p*** | **Sig** |
| --- | --- | --- | --- | --- | --- | --- |
| Treatment | NO_2_^-^ | | 1 | 0.135 | 0.27864 | n.s. |
|  | NO_3_^-^ | | 1 | 0.36 | 0.61 | n.s. |
|  | PO_4_^3-^ | | 1 | 0.002935 | 0.0597 | n.s. |
|  | SiO_2_ | | 1 | 0.9712 | 1.84E-12 | *** |
|  | dFe | | 1 | 51.94 | 0.00565 | ** |
| Time | NO_2_^-^ | | 6 | 3.06 | 0.00155 | ** |
|  | NO_3_^-^ | | 6 | 5.65 | 0.661 | n.s. |
|  | PO_4_^3-^ | | 5 | 0.0088113 | 0.0804 | n.s. |
|  | SiO_2_ | | 6 | 0.2864 | 0.000273 | *** |
|  | dFe | | 6 | 289.44 | 9.25E-06 | *** |
| Treatment*Time | NO_2_^-^ | | 5 | 0.097 | 0.97062 | n.s. |
|  | NO_3_^-^ | | 5 | 5.05 | 0.6 | n.s. |
|  | PO_4_^3-^ | |  |  |  |  |
|  | SiO_2_ | | 5 | 0.2612 | 0.00026 | *** |
|  | dFe | | 5 | 37.4 | 0.31495 | n.s. |
| Residuals | NO_2_^-^ | | 36 | 4.026 |  |  |
|  | NO_3_^-^ | | 37 | 50.67 |  |  |
|  | PO_4_^3-^ | | 13 | 0.00897 |  |  |
|  | SiO_2_ | | 32 | 0.2543 |  |  |
|  | dFe | | 43 | 263.04 | 263 |  |

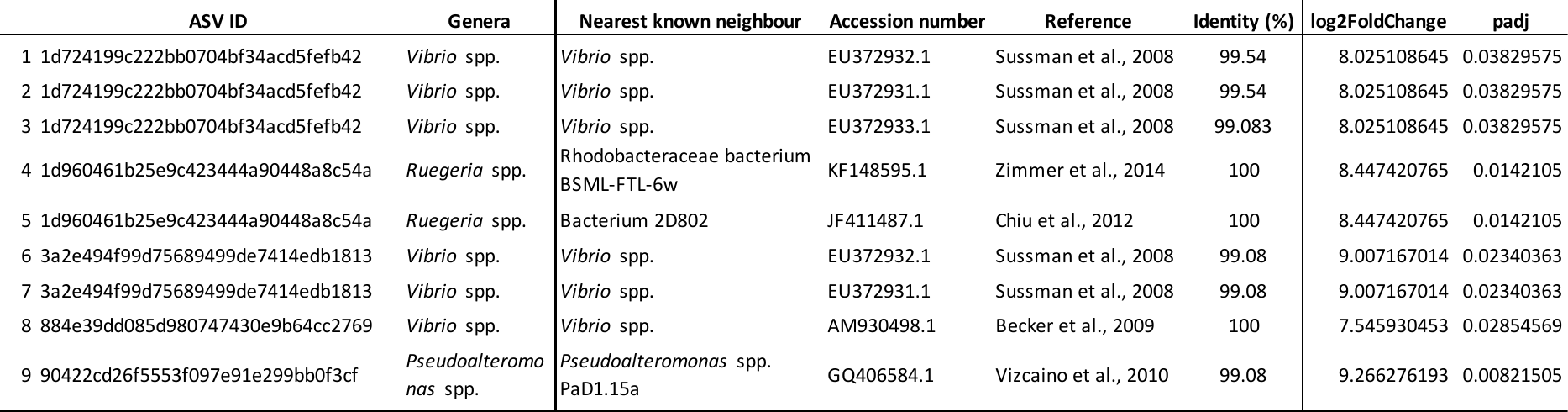

**Supplemental Table 5**. Log2 fold change for Amplicon Sequence Variants (ASVs) significantly enriched in storm samples with significant blast hits to known coral pathogens. DESeq function in the DESeq2 package was used. All E-values were equal to 0.

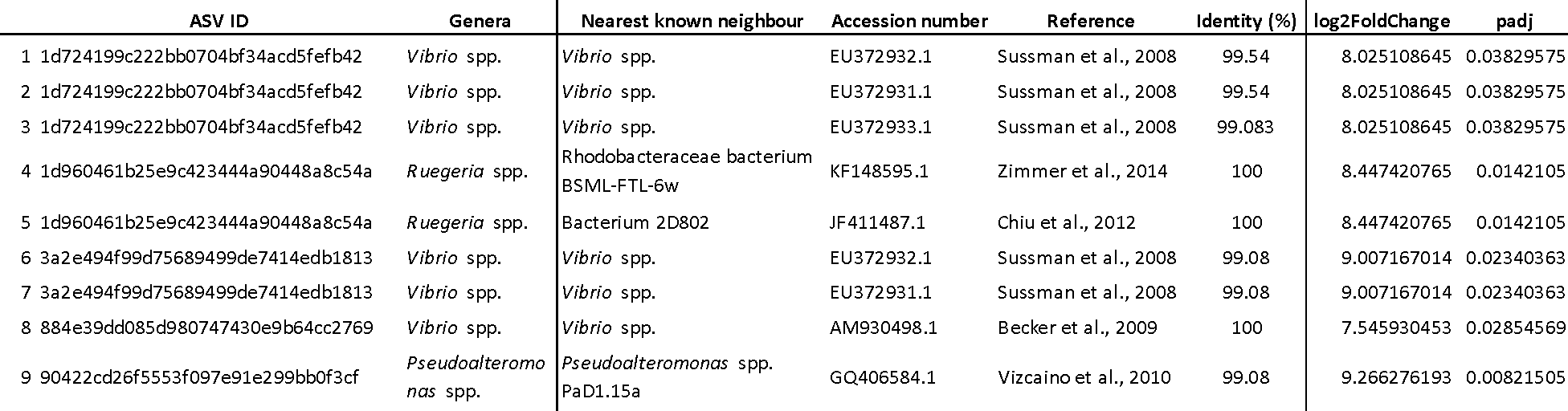
